# A human single-cell atlas of the Substantia nigra reveals novel cell-specific pathways associated with the genetic risk of Parkinson’s disease and neuropsychiatric disorders

**DOI:** 10.1101/2020.04.29.067587

**Authors:** Devika Agarwal, Cynthia Sandor, Viola Volpato, Tara Caffrey, Jimena Monzon-Sandoval, Rory Bowden, Javier Alegre-Abarrategui, Richard Wade-Martins, Caleb Webber

## Abstract

We describe a human single-nuclei transcriptomic atlas for the **Substantia nigra** (**SN)**, generated by sequencing ~ 17,000 nuclei from matched cortical and SN samples. We show that the common genetic risk **for Parkinson’s disease** (**PD**) is associated with **dopaminergic neuron** (**DaN**)-specific gene expression, including mitochondrial functioning, protein folding and ubiquitination pathways. We identify a distinct cell type association between PD risk and oligodendrocyte-specific gene expression. Unlike **Alzheimer’s disease** (**AD**), we find no association between PD risk and microglia or astrocytes, suggesting that neuroinflammation plays a less causal role in PD than AD. Beyond PD, we find associations between SN DaNs and GABAergic neuron gene expression patterns with multiple neuropsychiatric disorders. Nevertheless, we find that each neuropsychiatric disorder is associated with a distinct set of genes within that neuron type. This atlas guides our aetiological understanding by associating SN cell type expression profiles with specific disease risk.

## Introduction

The identification of the cell types relevant to a given disease is key to understanding the causal processes underlying the molecular aetiology. Many disorders, including neurological and psychiatric, are strongly influenced by genetic variation altering gene function. Therefore, by associating genetic variation with particular genes, those genes’ spatial expression patterns can then implicate specific cell types with that disorder. The cell type(s) associated with the genetic risk for a disease are not necessarily those cell types most directly associated with the defining symptoms. For **Alzheimer’s disease** (**AD**), intersecting risk-associated common genetic variation with cell type-specific gene expression proposed that the genetic risk for AD most significantly influences microglia, moving the research focus away from the neurons whose loss underlies symptoms, towards a role for neuroinflammation in this neuronal loss ^2^. It is thus critical to re-examine our aetiological assumptions when the data to do so are available.

The main hallmark of **Parkinson’s disease** (**PD**), the most common progressive neurodegenerative movement disorder, is the selective loss of **dopaminergic neurons** (**DaNs**) in the Substantia nigra (SN). Midbrain DaNs from the SN have a key role in the regulation of movement, cognition, motivation and reward, and their loss underlies deficits in fine motor control observed in PD^3^. However, while the loss of DaNs underlies key PD pathology, roles for other cell types such as astrocytes and microglia have been proposed that contribute to this loss ^4, 5^. Unfortunately, the human SN is sorely understudied compared with the neocortex, for which single-cell/nuclei studies ^6, 7^ have comprehensively mapped and characterised the diverse cell type populations and identified cell type-specific disease associations ^7, 8^. A comparable systematic and unbiased survey of cell-type-specific gene expression across the human SN will help us to identify the role of neuroglia in the selective vulnerability of the DaNs in PD, and provide significant insight into the potential contributions of the SN cell types to other neurological disorders.

In this study, single-nuclei RNA sequencing was performed on the human SN and cortex regions from the same individuals to resolve their regional cellular diversity and to identify region-specific neuronal and non-neuronal cell-type differences. From this cellular atlas, we mapped common risk variants for a range of brain-related disorders/traits to specific cell types. We performed a cell type-specific gene network analysis of PD and other psychiatric risk genes in the SN. Among many multiple disease risk/cell type association, we show that genetic risk in PD is indeed associated with DaNs, but also with oligodendrocytes, giving new insights into the causes of Parkinson’s disease and other disorders. By contrast, we find no association between PD risk and microglia, suggesting that neuroinflammation is not a key causal process in PD.

## Results

### Single nuclei sequencing of the human Substantia nigra and cortex

We sequenced the transcriptomic profiles of 10,706 nuclei and 5,943 nuclei from the cortex (middle frontal gyrus) and SN, respectively, of 12 matched samples (including 2 SN replicates) from five human postmortem brains using the 10x Genomics Chromium platform (**Supplementary Table 1**, **Supplementary Data 1**, **Methods**). We identified ten distinct cell populations across all samples within the SN (Figure 1a, **Supplementary Figure 1, Supplementary Note)**, which included (i) astrocytes with two subtypes: astrocyte-1 population was representative of the “A1” reactive neuro-inflammatory state, while the astrocyte-2 population aligns with the “A2” astrocytic state associated with growth and reparative functions^9^ (**Supplementary Table 2**)(ii) oligodendrocytes (ODCs), (iv) microglia cells, (v) oligodendrocyte precursor cells (OPCs) (vi) DaNs (*TH* & *SLC6A3*), neuronal population of the SN pars compacta (SNpc) **(Supplementary Figure 2)** and (vii) GABAergic neurons, neuronal population of the SN pars reticulata (SNpr) expressing gamma-aminobutyric acid (GABA) receptors *GABRA1* & *GABRB2* and the enzymes *GAD1* & *GAD2* required for GABA neurotransmitter synthesis (Figures 1a and 1c, Supplementary Data 2 and 3, Supplementary Figure 2). In the cortex, we identified six distinct cell populations including: astrocytes, **excitatory neurons** (**Ex**), **inhibitory neurons** (**In**), ODCs, OPCs and microglia, but no distinct cluster for endothelial cells (Figure 1b, **Supplementary Figure 3** and **Supplementary Figure 4**, **Supplementary data 2 & 3**, **Supplementary Note**). A joint clustering of both SN and cortex regions revealed that the cortex and SN form distinct clusters by cell types (neuronal cells; ODC; astrocyte, microglia, OPC and endothelial cells) rather than by region (**Supplementary Figure 1 d & e**).

**Figure 1:**
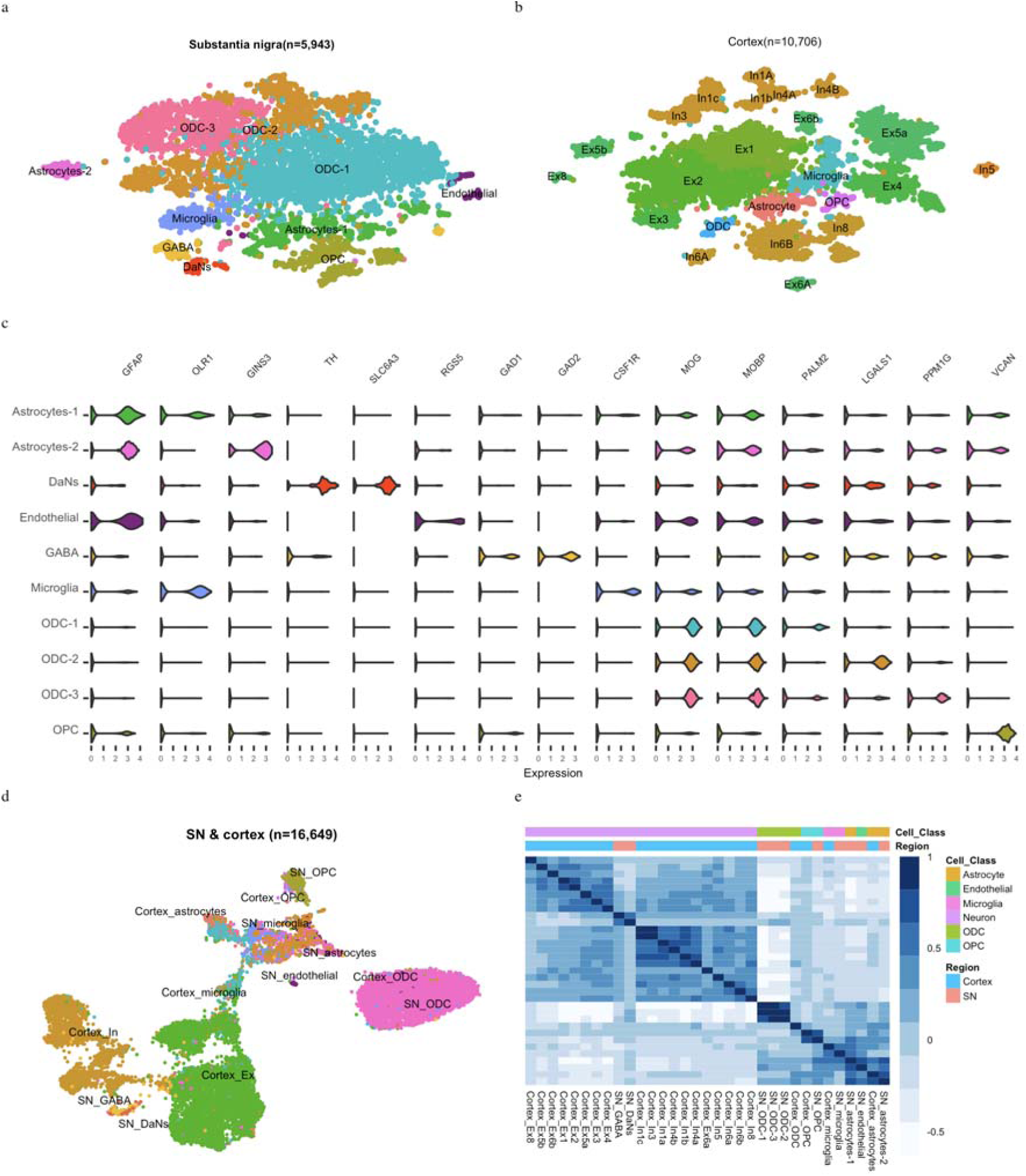
Single nuclei transcriptomic cell atlas of the adult human cortex and Substantia nigra (SN). T-distributed stochastic neighbour embedding (tSNE) plot of gene expression relationships amongst the single nuclei cells from 5 individuals in the (a) Substantia nigra (SN) (n=5,943) and (b) cortex (n=10,706). (c) Violin plots of expression values (log10 TPM values) of enriched cell type-specific markers for the cell types in the SN (Supplementary Data 3) (d) Uniform Manifold Approximation and Projection (UMAP) plot of both the cortex and Substantia nigra cell types (n=16,649 cells) showing distinct clustering by cell type. (e) Correlation heatmaps showing hierarchical clustering of Pearson correlation scores calculated between averaged cell type subclusters in both regions. The transcriptional correlation is largely explained by cell type and not by the region of origin. DaNs: Dopaminergic neurons; Ex: Excitatory neurons; GABA: GABAergic neurons; In: Inhibitory neurons, ODC: Oligodendrocytes; OPC: Oligo-precursor cells.

We captured a higher proportion of nuclei from glia in the SN (95.5% glia) represented mainly by ODCs (72%)(**Supplementary Table 3a**) than that obtained from the cortex (12% glia) (**Supplementary Table 3b**). As our cortex and SN atlases have been generated from the same individuals through the same process, this difference in glial proportion likely reflects genuine variation in the cellular composition of different brain regions. The observed proportions of different cell populations in our SN atlas were consistent with histopathological studies, reporting oligodendrocytes as the most frequent glial cell population (45-75% across all brain regions, 62% for SN)^10^. We observed consistent clustering by cell type between replicates, across samples and regions (**Supplementary Figure 5**), suggesting we have repeatedly captured the same resident cell populations.

### The single-cell profiles of the Substantia nigra inform on the cell types that the genetic risk of Parkinson’s disease and other disorders likely manifests within

We used the cell type-specific gene expression patterns from the human SN transcriptomic cellular atlas to identify specific SN cell types through which genetic variants contributing to each of 30 human complex traits (**Supplementary Table 4**) might be acting, employing two distinct methods (**LD score regression** (**LDSC**) ^11^ and **Multi-marker Analysis of GenoMic Annotation** (**MAGMA**) ^12^) (**Supplementary Table 5**). We observed for the first time in human a significant association between PD genetic risk and genes with DaN-specific expression patterns (q_*MAGMA*_=4.6×10^−3^, Figure 2a). We also identified a second association between PD genetic risk and genes with ODC-specific expression patterns (q_*MAGMA*_=0.035; Figure 2a). We demonstrated that the fraction of PD genetic risk contributing to the ODC association is distinct to that fraction of risk associated with DaNs (**Supplementary Table 6**, p=7×10^−3^), proposing distinct PD-associated cell etiologies within the SN. In the cortex map, we found a significant Ex neuron cell association with PD genetic risk variants (q_*MAGMA*_=6.1×10^−3^, Figure 2b, **Supplementary Table 7**). Our cortex/SN paired tissue sample study design enabled conditional analyses without effects due to individual/study variation to be performed for cell types across these two brain regions (**Supplementary Note**). We examined the effects of the differing neuron/glial cell proportions between the SN and cortex upon genetic risk/cell associations by creating artificially matched SN/cortex cellular atlases possessing the same proportion of glial cells by randomly sampling the original cellular populations (**Methods**). We observed a large agreement in the genetic risk/cell associations between the original and the homogenous cell atlas (**Supplementary Figure 6**) (SN R=0.83 (p < 10^−16^) & cortex R=0.72 (p < 10^−16^)), showing that the cellular proportions do not obscure the cell type-associations within the same tissue. Nevertheless, only the cross-tissue cell type conditional analysis conducted within the homogeneous cell-atlas revealed that the PD genetic risk associated with the SN DaNs and cortex Ex gene expression profiles was indistinguishable (**Supplementary Table 8**, p= 0.15 (homogenous cell atlas)/ p=0.049 (original cell atlas)). Concordantly, we observed that known PD risk genes^13^ are generally more highly expressed in neuronal cell types across the SN (**Supplementary Figure 7**).

**Figure 2:**
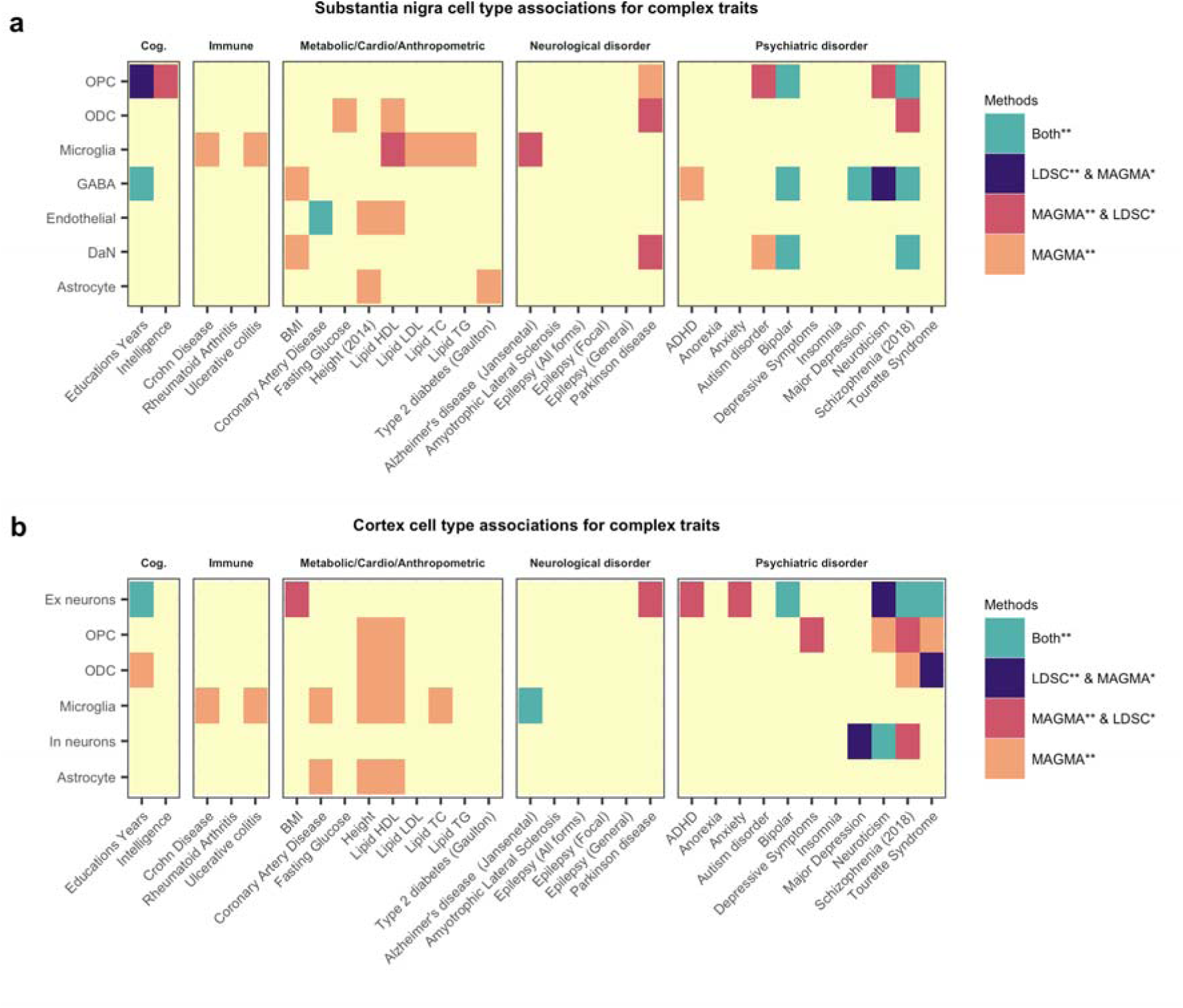
Identification of relevant brain Substantia nigra and cortex cell type involved in diverse human complex traits by using a single human nuclei transcriptomic atlas. We used two approaches to identify the associations between genetic risk variants of different complex trait and cell types from (a) Substantia nigra and (b) cortex: stratified LD score regression (LDSC) and the MAGMA gene set analysis. The heatmap colours give different degrees of significance with both methods or either method, * and ** indicate nominally significant p-value (< 0.05) and q value (correction for the number of cell types tested). The different traits were clustered by category: Cognitive phenotype (Cog.), Autoimmune disease (Immune), Metabolic, Cardiovascular and Anthropometric Traits (Metabolic/Cardio/Anthropometric) Neurological disorder, Psychiatric disorder.

Our SN atlas associates the genetic risk of neuropsychiatric disorders with DaN gene expression but also with GABAergic gene expression, e.g. schizophrenia (SCZ) (DaN and GABA) (Figure 2a). However, the conditional analyses between cell types of the SN demonstrated that where DaN and GABAergic neurons are both associated to a neuropsychiatric disorder, the association with DaNs is lost once conditioned upon GABA neuronal expression but not vice versa (**Supplementary Table 6**). Thus, the genetic risk of these psychiatric disorders is more broadly associated with genes expressed in the GABAergic neurons of the pars reticulata than the DaNs of the pars compacta. As previously reported^7^, we found in the cortex significant neuropsychiatric disorder cell type associations with both Ex and In neurons (e.g. SCZ) (**Supplementary Table 7)** and that these associations are most often distinct from SN neurons (conditional analysis; see **Supplementary Table 8**) proposing a distinct role of the SN in the aetiology of neuropsychiatric disorders, especially SCZ. Furthermore, we found a significant oligo-type association for different neuropsychiatric disorders (e.g. SCZ risk with two SN glial cell populations, namely OPC (q_LDSC_= 4.27×10^−3^,q_*MAGMA*_=1.32×10^−4^) and ODC (q_*MAGMA*_=1.36×10^−5^)) (**Figure 2a**). These associations support the hypothesis that for many brain disorders, glia may causally contribute to the neuronal alterations. This is well illustrated by the association of AD risk variants with microglia-expressed genes, found here for microglia in both the SN (q_*MAGMA*_=9.81×10^−4^) and the cortex (q_*LDSC*_= 0.01,q_*MAGMA*_=7.2×10^−4^) (Figure 2, **Supplementary Table 5** and **Supplementary Table 7**) and indistinguishable between microglia populations from the two brain regions (**Supplementary Table 8**, p=0.84).

There are significant overlaps in the genetic risk between different neuropsychiatric disorders^14^. We examined whether a common cell type association between two neuropsychiatric disorders was associated with a shared genetic risk between those disorders by re-performing the cell-association analysis using the genetic risk of one disorder after conditioning on the genetic risk effect of the other disorder. In general, these conditional analyses suggest that an overlapping component of risk between these disorders is associated with OPCs while distinct non-overlapping fractions of genetic risk act through the same neuronal types (Figure 3). In some cases we noted that a cell-specific association disappeared non-reciprocally when conditioning the risk on other disorders (e.g. GABAergic neurons and ADHD or Ex neurons and anxiety; Figure 3), reflecting a more restricted and subsumed cell-specific genetic risk association for that disorder.

**Figure 3:**
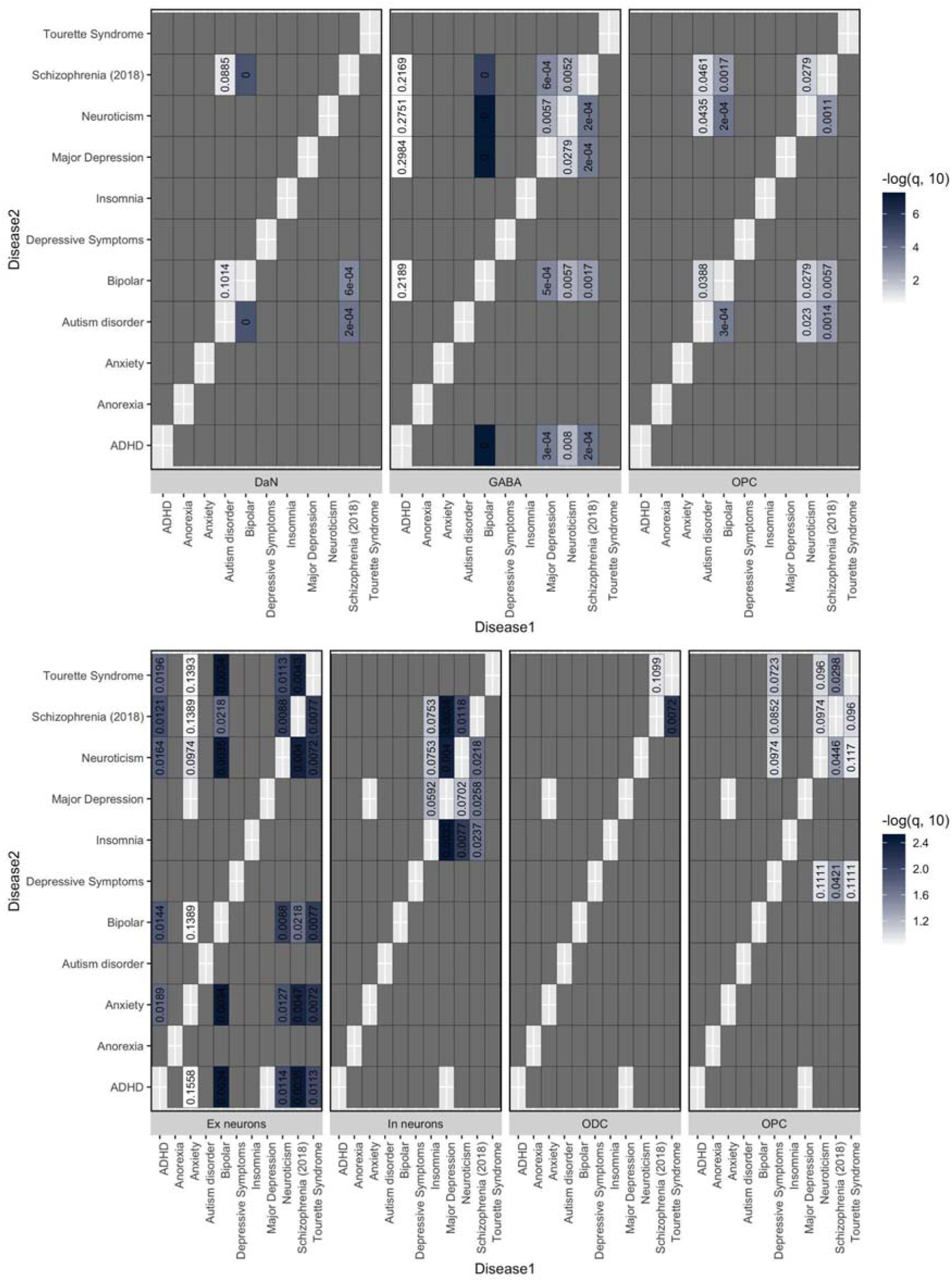
Evaluation of overlapping cell type-associations between any two neuropsychiatric disorders to identify a shared cell-type-specific component of risk, for the SN cell types (top) and the cortex (bottom). Each heatmap represents the results from LDSC of the associations of a specific cell type expression profile with the genetic risk of a given neuropsychiatric disorder (Disease1-X-axis) after conditioning on the genetic risk of another neuropsychiatric disorder (Disease2-Y-axis). This analysis was only performed where two neuropsychiatric disorders showed a significant (or suggestive) association with the same cell type (Figure 2). The blue heatmap colours are proportional to -log 10 q-value (FDR-adjusted p-value) of the enrichment of genetic variants associated with a disorder adjusted for another disorder. The cell-associations that were not evaluated are coloured in dark grey.

Finally, we also found genes specifically expressed in SN cell types to be associated with traits not specific to the brain. We identified a unique cell type-specific signal between microglia and genetic variants associated with High-density lipoprotein (HDL) cholesterol level (q_*MAGMA*_= 0.046)(Figure 2a). Given the role of lipids in AD ^15, 16^, we asked whether the HDL cholesterol association was related to the association between AD risk and microglia gene expression (Figure 2a). However, conditional analysis suggests distinct genetics underlie the associations of AD and HDL cholesterol levels with microglia (**Supplementary Figure 8**).

### Genetic risk highlights cell-type-specific networks and pathways

Cell type-specific gene expression patterns within the SN and cortex may help identify cellular circuitry that underlies disease-associated cell-type-specific vulnerabilities. We used protein-protein interactions (PPI) as evidence for functional relationships between genes, and built cell type-specific PPI networks within which we identified a number of modules of highly interconnected genes (**Methods** and **Supplementary Note**). For each significant cell type/disease-risk association discovered above (Figure 2), we looked to refine the disease-risk association to a relevant cell type-specific gene module using MAGMA gene set analysis (Figure 4). PD-risk showed association to nigral DaN modules (M1 and M2) (Figure 4b, **Supplementary Figure 9** and **Supplementary Figure 10**) enacting processes previously associated with PD such as mitochondrial organisation and functioning, endocytosis, protein ubiquitination and macroautophagy ^17^. PD-risk association to ODCs (M5) was related to genes enriched in metabolic processes, gene regulation, kinase activity, protein phosphorylation and neurogenesis while the OPC risk-associated module (M1) was enriched in metabolic processes, gene regulation and cell differentiation. The SCZ-risk association to cortical neuronal populations was mainly related to synaptic signalling and neuronal developmental processes (Ex neuron module M1/In neuron M1 and Ex neuron module M4/In neuron M4, respectively; Figure 4a and **Supplementary Figure 9**). In the nigral neuronal population, genes within DaN module M4 and GABA module M3, both associated with synaptic signalling/neuron development, were also enriched in SCZ-risk (Figure 4b and **Supplementary Figure 9**). A further SCZ association to DaN (M1) was found related to mitochondria (Figure 4b), supporting the role of mitochondrial dysfunction in SCZ ^18^. Lipid metabolism and neuron development were found enriched in both cortical ODCs (M2) and OPCs (M4) modules associated to SCZ-risk. Furthermore, nigral OPC modules (M3 and M5), related to nervous system development and synaptic signalling, were found associated with SCZ-risk (Figures 4b and **Supplementary Figure 9**). The nigral OPC M3 module also showed a significant association with other traits such as neuroticism (corrected p-value = 0.004). As predicted by the genetic risk, a BP association was found to the nigral DaN module (M4) associated with synaptic signalling and neuronal development (Figure 4b and **Supplementary Figure 9**).

**Figure 4:**
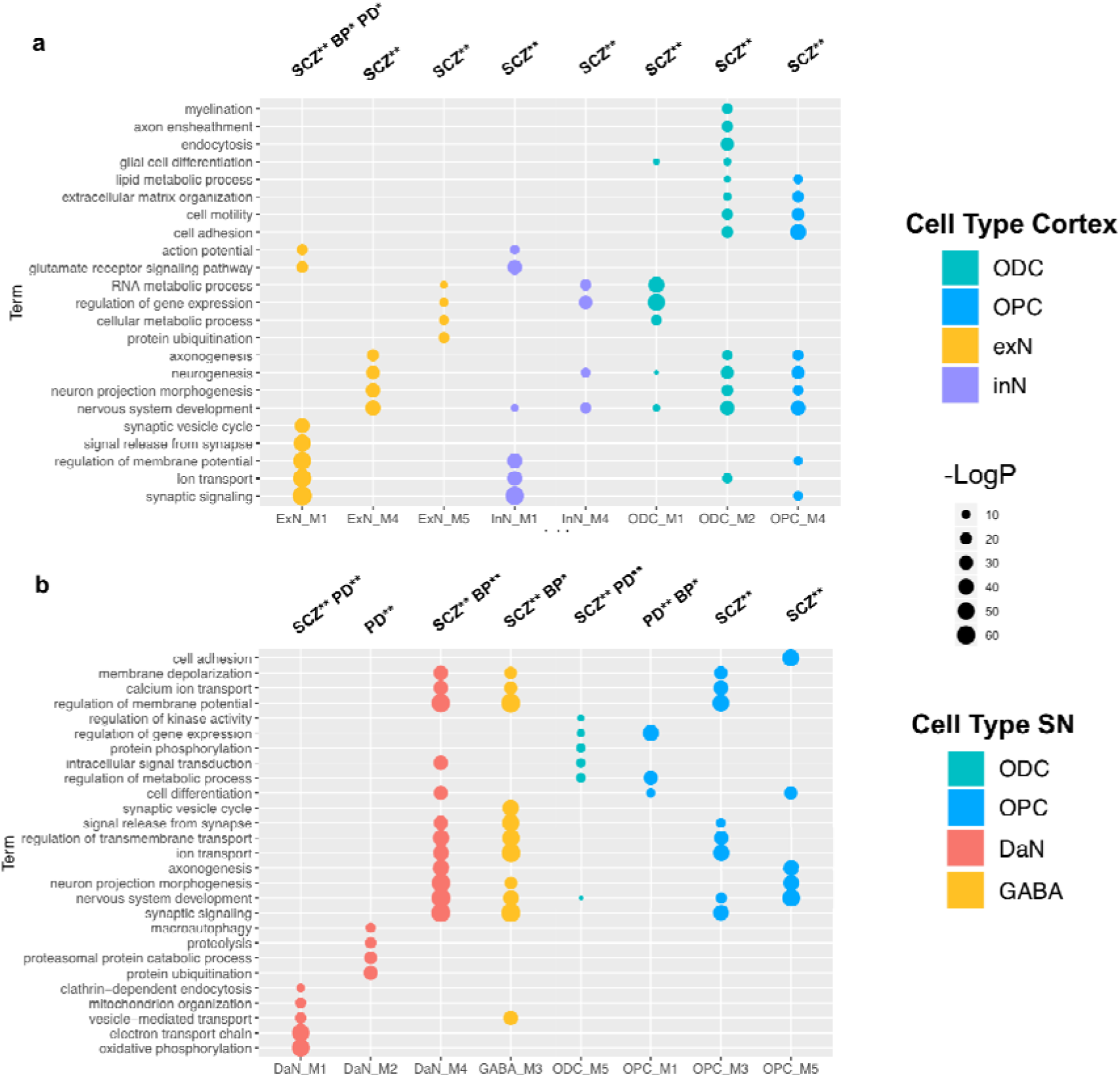
GO enrichment for cortex (a) and SN (b) cell type-specific gene modules enriched in PD, SCZ and BP -risk. We tested the convergence of disease genetic risk at a functional level across cell-type-specific PPI gene modules. The top representative Gene Ontology (GO) Biological Process (BP) terms are shown for cell type modules with either PD, SCZ or BP risk enrichment that reached significance in the more general cell type level analysis (Figure 2). Tests across all identified gene modules are reported in Supplementary Figure 10, and complete lists of enriched GO BP terms are reported in Supplementary Table10 and S11. The size of circles represents -log(p-value) for GO enrichment; colours correspond to cell types; single star (*) corresponds to nominally significant p-values, and double stars (**) correspond to multiple tests corrected significant p-values.

We then evaluated whether a common module association within the same cell type between two disorders was associated with a shared genetic risk between those disorders. For this, we re-performed the cell-specific module-association analysis using the genetic risk of one disorder after conditioning on the genetic risk effect of the other disorder and vice versa. For the three nigral modules associated with two disorders (Figure 4), we found the associations of PD and SCZ to DaN module M1 were independent (p-value = 0.0002 for PD conditioned on SCZ and p-value = 0.0009 for SCZ conditioned on PD) with PD-associated M1 genes mainly enriched in endocytosis processes (*regulation of endocytosis*, GO:0030100 p-value = 0.0003) and SCZ-associated M1 genes mainly enriched in neuronal migration and development (*neuron migration*, GO:0001764 p-value = 0.0004). Similarly, we found the associations between BP and SCZ to DaN M4 to be independent (p-value = 0.0003 for BP conditioned on SCZ and p-value = 0.002 for SCZ conditioned on BP) with BP-associated M4 genes mainly enriched in synaptic transmission processes (*synaptic signaling*, GO:0099536 p-value = 3.9e-= and SCZ-associated M4 genes mainly enriched in neuronal development (*nervous system development*, GO:0007399 p-value = 6.3e-06). By contrast, we found that association of the nigral ODC M5 module with PD risk was lost when conditioned on SCZ risk (p-value = not significant) but that the SCZ association remained when conditioned on PD risk (p-value = 0.002) revealing convergence, specifically that the SCZ risk loci overlapped and subsumed the loci underlying the ODC M5 PD association.

## Discussion

We generated a comprehensive human single-nuclei transcriptomic atlas for SN by sequencing ~ 17,000 nuclei from matched cortical and SN tissue enabling the identification and comparison of brain region-specific cell types. In this study, (i) we re-identify all the major cell types previously reported indicating sufficient genes/cells coverage, (ii) we find a large difference in the neuronal-glial cell composition between cortex and SN that confounds bulk tissue disease associations (Figure 1, **Supplementary Figure 11** and **Supplementary Table 9**), (iii) we identify multiple associations between the genetic risk of particular diseases and specific cell types in the nigra and cortex (Figure 2), (iv) we find that where there are multiple neuropsychiatric disease associations for a given neuron type, the loci associated for each disorder are distinct and do not appear to converge on the same set of genes within that cell type, while for glia we do observe convergence (Figure 3) and (v) we determine the different cell-specific gene-networks and their functions perturbed by disease risk variants (Figure 4).

This SN/cortex paired-sample atlas is a valuable resource to interpret the genetic architecture of many disorders, especially for PD and other disorders with particular vulnerabilities in the SN as compared to the cortex. However, the fraction of PD genetic risk that mapped to cortical Ex neurons was indistinguishable from that mapping to SN DaNs, and thus our atlas alone is unable to propose why SN DaNs is lost earlier in PD pathology. Our cellular atlas suggests similar cellular functions associated with PD genetic risk across neuronal cell types in PD nigral and cortical degeneration^3^. Thus, different metabolic demands and different local environments of these neurons from different brain regions likely contribute to their distinct vulnerabilities ^19^. PD genetic risk also appears to manifest through ODCs and OPCs (Figure 2), which implicate gene regulation in metabolic processes and in cell development and support the growing evidence of the role of glia in neurodegenerative disorders ^20^. Notably, we observe expression of the known PD gene *LRRK2* to be significantly higher in OPCs than other SN cell types (**Supplementary Figure 7**). A role for ODCs in PD is surprising given the light myelination of SN DaN axons^21^, but our relatively unbiased approach allows for unexpected associations that require further understanding. Indeed, multiple studies have identified white matter impairments, which correlate with progression and appear to precede grey matter atrophy, and our results may provide a missing molecular link to PD genetic risk ^22^. Unlike AD, we do not find an association between PD genetic risk and microglia, suggesting neuroinflammation may play a lesser role in PD risk than in AD risk.

The Midbrain encompassing the SN is a key brain region of interest after the cortex for several neuropsychiatric disorders ^23^. For SCZ we found distinct associations with both nigral neurons and cortical neurons. However, it may be that this SCZ-DaN association relates to DaNs within the adjacent **Ventral tegmental area** (**VTA**) region previously implicated in several neuropsychiatric disorders ^24^. Without corresponding VTA atlases from these brains, we are unable to perform the relevant conditional tests, which highlights the value of larger future studies capturing more regions from the same brains. Due to the key role of DaNs in PD, the role of GABAergic neurons of the pars reticulata in other diseases is relatively understudied. Nonetheless, the GABAergic neurons project to the prefrontal cortex and nucleus through mesolimbic pathway and control mainly the reward system (learning about motivationally relevant stimuli in the environment), which is affected in patients with neuropsychiatric disorders ^25^ and our novel associations suggest these neurons have the potential to be influenced by the genetic risk of several neuropsychiatric disorders. Across several neuropsychiatric and neurodegenerative disorders, we found multiple glia association, especially to OPC processes related to synaptic signalling ^26^.

We describe here the first comprehensive human SN cell type atlas. Together with a matching cortical atlas, these atlases allow the systematic characterisation of SN cell types and the identification of cell type-specific processes influenced by multiple complex disorders.

## Online Methods

### Samples

Five controls were selected on the basis of the absence of neurological clinical disease and by midbrain **RNA integrity number** (**RIN**) yielding scores over eight from the Oxford Brain Bank (**Supplementary Table 1**). However, the histopathological examination revealed **cerebral amyloid angiopathy** (**CAA**) in the brain of one individual (Sample 3, **Supplementary Table 1**). CAA is one of the morphologic hallmarks of Alzheimer disease (AD) ^27^, but it is also, very common in the brains of elderly patients who are neurologically healthy ^28^. To ensure that this individual did not affect the results and conclusions of this study, we repeated all further analyses with and without the samples from this individual and found a very high correlation between results with highly concordant cell clustering (see **Supplementary Note**) and thus retained this sample for greater power. Informed consent had been collected from all cases fulfilling the requirements of the Human Tissue Act 2004. Blocks of approximately 3 mm^2^ were dissected from snap-frozen slices by a qualified neuropathologist (J.A-A) from the central portion of the Substantia nigra at the level of the third nerve encompassing both ventral and dorsal tiers and from the middle frontal gyrus encompassing the cortex but macroscopically excluding the subcortical white matter. Replicate blocks obtained on a different day were obtained in two out of the five individuals.

### 10x sequencing data

10x Chromium single nuclei sequencing was performed on the cortex and SN regions from five individuals. In total 12 samples (including 2 SN replicates), with a total of 12,015 and 6,105 nuclei were sequenced for the cortex and SN over two days (**Supplementary Table 1**). Reads were processed and mapped to the Human Genome (GRCh38.84-premrna) with Cell Ranger 2.1.1 (**Supplementary Note**). We recovered a median number of 2,455 and 690 nuclei after sequencing for the cortex and SN. The multiple sample libraries were sequenced with mean reads ranging from 46,598 - 59,513 and 18,377-44,710 for the cortex and SN, respectively. Overall we detected median genes per nuclei ranging from 607-3,364 for the multiple individuals across both regions (**Supplementary Data 1**). Similar in range as other human brain single nuclei studies ^7, 29^, the mean read depth per nuclei was 43,150 (95% CI: 34,488-51,821) and the median number of genes detected per nucleus was 1,886 (95% CI:1,228-2,544) **(Supplementary Data 1)**. For the SN single nuclei atlas, we generated biological replicates and showed that cell profiles from the same individual were comparable to each other (**Supplementary Figure 12, Supplementary Note**).

### Data processing

The filtered **Unique Molecular Identifiers** (**UMI**) feature-barcode matrices generated with CellRanger 2.1.1, were processed with Seurat R package (v2.3.4)^30^, separately for the SN and the Cortex. As quality control steps we retained genes with a count of 1 in at least 3 nuclei and removed nuclei with < 500 genes per sample, high thresholds of nUMI (range of > 5000-12000, sample dependent), mitochondrial percentage > 0.05 and ribosomal percentage > 0.05. The distributions of number of genes, number of UMIs and percentage of reads mapped to mitochondrial and ribosomal genome were further inspected for quality assurance. Each normalised sample was linearly regressed to remove any inter-cellular gene expression variation because of technical effects associated with UMI coverage, ribosomal and mitochondrial percentage, followed by gene level scaling of the data by using the “ScaleData” function in Seurat. Cell cycle phase scores were predicted for each cell per sample and determined to not be an important source of variation and bias in the SN and cortex (**Supplementary Figures 1 & 3**).

### Cell clustering analysis

After merging the samples, identifying common sources of variation based on the **highly variable genes** (**HVG**) (**Supplementary Note**), performing a **canonical correlation analysis** (**CCA**)^30^, and discarding rare non-overlapping cells between samples, 5,943 nuclei and 22,736 genes (15,568 protein coding genes) remained for the SN and 10,706 nuclei and 26,145 genes (16,423 protein-coding genes) remained for the cortex. The CCA analysis identified the top numbers of CCA vectors to align for the SN and cortex as 25 and 42 dimensions. The shared-nearest neighbour graph was constructed on a cell-to-cell distance matrix from the top aligned CCA vectors followed by Louvain clustering^31^ to identify cell-type clusters, which were visualised with **t-distributed stochastic neighbour embedding** (**t-SNE**) and **Uniform manifold approximation and projection** (**uMAP**) plots.

On the basis of previous knowledge, consistency and validity of the different resolutions, we selected the final number of clusters based on the clustering resolution for the Louvain algorithm in the range of 0.4 −0.8, which included all the major cell-types and subtypes (**Supplementary Note**). Cell clusters with fewer than 30 cells were omitted from further analysis.

### Cell type annotation for cortex and SN

We characterised the cellular identities of clusters in the SN (**Supplementary Note**, Figure 1a,c) and the cortex (**Supplementary Note**, Figure 1b) nuclei by identifying known marker genes enriched in each of the clusters (**Supplementary Note**)^7, 32^. For each of the sub-clusters, the enriched marker genes were identified by differential expression of the cells grouped in each sub-cluster against the remaining cells within the corresponding broad cell type cluster. This process resulted in the annotation of 23 and 10 different neuronal and non-neuronal cell types for the cortex and SN, respectively (Figure 1a,b).

### Differential Expression

Differentially expressed genes between cell types and subtypes were identified within Seurat^30^ by using the negative binomial test (false discovery rate (FDR)-corrected p-value <0.05) to identify 0.25 log fold enriched genes detected in at least 25% of cells in the cluster of interest. Differential expression analyses were performed for all clusters separately in the SN and cortex, for SN astrocyte and DaN subtypes and for cortical Ex and In neuronal subtypes (**Supplementary Data 3**).

### Comparison of 10x data sn-RNAseq data with published data

Comparison of the cortex and SN nuclei averaged cell-type populations with previously published single cell human temporal cortex (TC)^7^, single nuclei ^6^ and bulk SN laser-capture microscopy (LCM) DaNs ^8^and glial cell-types^9^ data further confirmed the broad cell-type annotations and, in the case of cortex, neuronal subtype classification consistency (**Supplementary Figure 13, Supplementary Note**). In particular, SN DaNs identified in our 10x study cluster with external LCM post-mortem DaNs (**Supplementary Figure 13**) and astrocyte-1 or reactive astrocyte population cluster with the external LCM astrocyte and microglia samples (**Supplementary Figure 13**). In addition, cell-types are comparable between single nuclei and single cell data (**Supplementary Figure 13**). A correlation > 0.85 was observed between all the major cortical cell-types and layer-specific neuronal subtypes between single nuclei cell atlas interrogated in this study and the human visual cortex (VC) and frontal cortex (FC) single nuclei-drop seq data ^6^ (**Supplementary Figure 13**). The data processing and method of comparison for each of the external datasets is described in Supplementary Information.

### Cell type association analysis

We intersected cell type-specific expression patterns with genetic risk of specified disease to identify disease-relevant cell types in SN and cortex for 30 diseases and traits (Figure 2, **Supplementary Table 4**, **Supplementary Note**). We performed these cell type association analyses with two commonly used approaches: LDSC^11^ and MAGMA^12^. While these methods are complementary, we confirmed that their findings were well aligned (**Supplementary Figure 14, Supplementary Note**). Where we identified multiple cell types associated with the same trait, we performed conditional analyses to evaluate whether it was the same set of genetic variants acting in different cell types or distinct sets of genetic variants in each cell type suggesting multiple cellular aetiologies. Where traits/disorders were associated with both SN and cortical cell types, we considered that the different proportions of glial versus neurons for these two brain regions could bias the identification of cell-type-specific genes. To test this potential bias, we repeated the conditional analysis with artificially homogeneous SN/cortex cellular atlas, which included the same proportion of glial cells (**Supplementary Note**). To evaluate the genetic overlap between two traits, each showing an association with the same cell type, we performed GWA conditional analysis with **multi-trait-based conditional & joint analysis** (**mtCOJO**)^33^ (Figure 3). We then repeated LDSC analyses with the GWA summary statistic of one trait adjusted for the second trait and vice versa.

### Functional Analysis

#### Cell type-specific protein-protein interaction (PPI) network and identification of gene modules

A cell type-specific ***protein-protein interaction*** **(PPI)** network is built by extracting PPIs from PPI network between cell type-specific genes (**Supplementary Data 4**). To identify modules of highly interconnected genes in a cell type-specific PPI network, we employed “cluster_louvain” function in “igraph” R package ^34^. This function implements the multi-level modularity optimization algorithm ^35^, where at each step genes are re-assigned to modules in a greedy way and the process stops when the modularity does not increase in a successive step. Modules with > 30 genes are used for further analysis as smaller modules have low informativity. MAGMA gene set analysis was used to test enrichment in disease-risks across all identified modules.

#### Gene Ontology pathway enrichment analysis

We performed gene ontology (GO) enrichment analysis with topGO ^1^ R by testing the over-representation of gene ontology biological processes (GO BP) terms within the input gene sets using Fisher test. Revigo was used to summarise the top 100 enriched GO BP terms in a smaller number of categories.

### Data sharing

The processed 10x 3’ Chromium single nuclei RNAseq UMI-barcode matrices for each sample are available from the Gene Expression Omnibus under the accession code GSE140231.

## Acknowledgements

CW is supported by the UK Dementia Research Institute funded by the Medical Research Council (MRC), Alzheimer’s Society and Alzheimer’s Research UK. CS is supported by the Ser Cymru II programme which is part-funded by Cardiff University and the European Regional Development Fund through the Welsh Government.

We acknowledge the Oxford Brain Bank, supported by the Medical Research Council (MRC), Brains for Dementia Research (BDR) (Alzheimer Society and Alzheimer Research UK), and the NIHR Oxford Biomedical Research Centre. The views expressed are those of the authors and not necessarily those of the NHS, the NIHR or the Department of Health.

Computation used the Oxford Biomedical Research Computing (BMRC) facility, a joint development between the Wellcome Centre for Human Genetics and the Big Data Institute supported by Health Data Research UK and the NIHR Oxford Biomedical Research Centre. The views expressed are those of the author(s) and not necessarily those of the NHS, the NIHR or the Department of Health. This work was supported by the Wellcome Trust via core funding to the Wellcome Centre for Human Genetics (award 203141/Z/16/Z). R W-M, J A-A and T C were supported by the Monument Trust Discovery Award from Parkinson’s UK.

## Contributors

DA, CS, VP performed data analyses and wrote paper, TC developed and performed nuclei extraction protocols, J M-S performed data analyses, RB - Performed sequencing and generated data, contributed to experimental design, J A-A contributed to experimental design and the selection and sourcing of post-mortem tissue, R W-M developed nuclei extraction protocols and contributed to experimental design, CW– conceived project, designed experiments, designed and led analyses, wrote paper.

