## Supplemental Material for "A human single-cell atlas of the Substantia nigra reveals novel cell-specific pathways associated with the genetic risk of Parkinson’s disease and neuropsychiatric disorders"

**Supplementary Note**

**Nuclei extraction**

Nuclei extraction was based on the protocol described by Krishnaswami *et al ^1^* with some modifications. Briefly, 3mm^2^ pieces of fresh-frozen brain tissue were placed in homogenization buffer (250 mM sucrose, 25 mM KCl, 5 mM MgCl_2,_ 10 mM Tris Buffer pH 8.0, 1 mM DTT, 1X protease inhibitor, 0.4 U ul^-1^ RNaseIN, 0.2 U ul^-1^ Superasin, 0.1% v/v Triton X-100) for 10 minutes prior to mechanical disruption with a dounce homogenizer. Homogenate was filtered through a 35 mM cell strainer before concentrating nuclei by centrifugation 400 g 5 minutes. Nuclei were resuspended in FACS buffer (1 XPBS, 1 RNase-Free BSA, 0.2 U μl^−1^ of RNasin Plus RNase inhibitor, 10 ng ml^-1^ Hoechst 33342) prior to FACS to obtain a population of nuclei free cellular debris.

**Single nuclei processing and 10x genomics library preparation**

Nuclei were washed once in PBS and resuspended in 0.04% BSA in PBS for loading on the 10x Chromium 3’ chip. Cell capturing, and library preparation was carried as per kit instructions (Chromium Single Cell Kit [v2 chemistry]). 3000 nuclei were targeted for capture per sample, after cDNA synthesis, 12-14 cycles were used for library amplification. The resultant libraries were size selected, pooled and sequenced using 75bp paired-end sequencing protocol on an Illumina HiSeq 4000 instrument. We used 12 samples for the 10X emulsions, with matched cortex and SN samples from 5 individuals with biological replicates for **substantia nigra** (**SN**) sample 4 (N4, N4B) and 5 (N5, N5B) run on two separate days (**Supplementary Table1**).

**10x sequencing data processing**

Reads were processed and mapped to the Human Genome (GRCh38.84-premrna) with Cell Ranger 2.1.1. We used default mapping arguments for cellranger “count” function to merge multiple sequencing runs for the same sample libraries. In order to have counts from both unspliced/pre-mRNAs and mature mRNAs, a custom human (GRCh38.84) “pre-mRNA” reference was created, whereby each gene transcript was annotated as an exon, so that pre-mRNA intronic reads can also be included in the UMI counts for each gene and barcode. The filtered gene /barcode matrices containing only the detected cell-associated barcodes and UMIs produced as a result were then further analysed.

**Dimensionality reduction and cell clustering analysis**

Seurat R package (v2.3.4) was used to integrate different samples for the cortex and SN and then to perform a dimensionality reduction analysis. **Highly variable genes** (**HVGs**) were identified from a mean variability plot (average expression versus dispersion (variance/mean) assigned to 20 bins based on average expression) using a log(variance/mean) cutoff of 0.5 and maximum cut-off of 3.5 to identify outliers for each sample. Within each bin, a z-score of log-transformed dispersion measure (variance/mean) was calculated. A z-score cutoff of 0.5 was applied to identify the highly variable genes for each sample and the union of top 3000 of these from each sample resulted in 1031 and 2243 genes in the SN and cortex, respectively. A **canonical correlation analysis** (**CCA**)^2^ was performed and after discarding rare – non-overlapping cells, the top numbers of CCA dimensions to align were identified by examining the correlation strength of all vectors through the biweight midcorrelation, a median based similarity metric saturation plot. This enabled to identify 25 and 42 canonical correlation vector dimensions to align for the SN and cortex.

The shared-nearest neighbour graph was constructed on a cell-to-cell distance matrix firstly by calculating the k-nearest neighbours with k=30 from the aligned CCA vectors. These were then used as input to the Louvain algorithm^3^ with different clustering resolutions for the SN and cortex to identify cell-type clusters. We performed the analysis with different resolutions in Seurat by sequentially increasing the resolution from 0.4 to 2.4 and assessing the clusters by constructing a phylogenetic tree for the averaged cluster populations at each resolution based on the HVG genes. For each branch of the subsequent tree a random forest classification error was calculated and an accuracy threshold of 80% was used to assess tree splits and cluster validity. This resulted in an optimal resolution parameter for the SN and cortex to be 0.4 and 0.8, respectively.

**Joint clustering analysis of SN and cortex samples**

Similarly using Seurat^2^, a joint clustering of the CCA aligned cortex and SN nuclei was performed using the **transcripts per million** (**TPM**) expression matrix of only protein-coding genes which were log-transformed and linearly regressed for nUMI. As before we used the “FindVariableGenes” function, a minimum log(variance/mean) cutoff of 0.5 and maximum cut-off of 9 on the mean-variability plot to detect outliers and identify 4221 HVGs. A **principal component analysis** (**PCA**) was then performed across all nuclei only using the HVGs. The standard deviations of the PCs were plotted on to a scree plot to identify the first 25 PCs as significant that were used as input to identify cell clusters by the **shared nearest neighbor** (**SNN**) based graph approach at a resolution of 0.4 through the “FindClusters” function in Seurat. The identified clusters were then visualised as a **Uniform Manifold Approximation and Projection** (**UMAP**) plot with significant PCAs as input to understand the global relationship between similar cell-types across the cortex and SN (**Figure 1d**). We found that changing the number of significant PCs in the range of 15-30 and the clustering resolution in the range of 0.4-0.8 did not identify very different clusters.

**Single-cell atlas of the substantia nigra**

All cell-types were mostly contributed to by all samples (**Supplementary** **Figure 1, Supplementary Figure 5**). Some sample-specific clustering and thus batch-specific effect can also be seen for reactive astrocyte-2 (N2B only, Day2) & ODC-3 (predominantly N2B) and this may be associated with heterogeneity between samples (**Supplementary Figure 2**, **Supplementary Table3** and **Supplementary Table1**). We found that neuronal populations have a higher UMI depth (median nUMI: DaNs = 8,419; GABA neurons =6,512) and gene coverage range (median nGene: DaNs= 3,579 ; GABA neurons= 3,529) than non-neuronal populations (median nUMI= 2,822;median nGene = 1,863)(**Supplementary Figure 1**).

**Single-cell atlas of the cortex**

The cortex cell-types were broadly classified into the major cell groups with the help of enriched genes marker and we resolved clusters as astrocytes (*AQP4^+^, GFAP^+^*), **Excitatory neuron**s (**Ex**)(*SATB2^+^,SLC17A7^+^*), Inhibitory neurons (In) (*GAD1^+^,GAD2^+^*) (see below), microglia(*CSF3R^+^,PLXDC2^+^*), oligodendrocytes cells (ODCs) (*MOG^+^,MOBP^+^*) and oligo precursor cells (OPCs) (*OLIG1^+^,VCAN^+^*) (**Supplementary** **Figure 4**, **Supplementary** **Data 3**). Similar to the SN, the major cell-types were contributed to by all samples, except for ODCs for which some sample-specific clustering (C1B & C3) can be seen (**Supplementary** **Figure 3**, **Supplementary Table 3**). Moreover, a uniform UMI depth, number of genes and cell-cycle phase trends were also, observed across the cortex for most cell-types, except for the Ex neurons (median nGene= 3,559; nUMI=7,332) and microglia (median nGene= 3,759, nUMI=7,330) showing higher UMI depth and gene coverage than other cell-types (median nGene = 2,096; nUMI=3,599) (**Supplementary Figure 3**).

**Cortex neuronal subtypes**

Lake e*t al ^5, 6^* have previously identified subtypes of Ex and In neurons based on specific markers that we used here to further, identify nine subtypes of Ex neurons and ten subtypes of In neurons drawn from across different cortical layers.

Neuronal subtypes were resolved into Ex and In neurons based on known marker genes *SATB2, SLC17A7, GAD1 and GAD2* (**Supplementary** **Figure 4**, **Supplementary Data 3**). The Ex neurons could be further classified based on their layer positions and known marker genes. A population of Ex2 subtype expressing the L2/3/ *CUX2* & L4 specific (*RORB^+^* ) markers and *COL5A2^+^* was the most prevalent in our dataset (19%, **Supplementary Table 3**), followed by the second most prevalent, L2-3 subpopulation (*LAMP5^+^*), Ex1 (14%, **Supplementary Table 3**) showing *CBLN2^+^,* *CUX2^+^* expression. While we did identify layer 4 subpopulations, Ex3 (*RORB^+^, COL5A2^-–^,IL1RAPL2^+^)* and Ex4 (*RORB^+^,TPBG^+^* ) like Lake et al^6^, (**Supplementary** **Figure 4**), we were not able to further distinctly resolve this sub-population into Ex3 (a-d) subtypes. We also, identified distinct subpopulations located in cortical L5*(PCP4^+^, ETV1^+^*), Ex5a (*PCP4^+^,HS3ST5^-–^*), Ex5b (*PCP4^-–^,HS3ST5^+^*) and Ex6a ( *HTR2C^+^,PCP4^+,^TLE4^+^*), with the last Ex6b (*HTR2C^-–^*, *TLE4^+^*,*SEMA3D^+^*) bordering on cortical L6 /6b or white matter(*NR4A2^+^*) (Figure s5). Lastly, the least prevalent (0.59%) cortical neuronal subpopulation we identified was Ex8 (*TLE4^+^,NTNG2^+^)* localised to L6.

In neuron, the subtypes were discriminated from the other cell-types by expression of *GAD1* and *GAD2* marker genes (**Supplementary** **Figure 4**) and further resolved into 10 subpopulations based on enriched marker gene expression that showed distinct profiles of established In neuron marker genes (**Supplementary Data 3**). We resolved spatially distinct *CNR1* expressing L1-2 subpopulations as In1a (*RELN^+^*), In1b (*THSD7B^+^*), In1c ( *VIP^+^,TAC3^+^)* and In3 (*VIP^+^,TSHZ2^+^*), but not the upper layer In1d and In2 subtypes (**Supplementary** **Figure 4**). We further identified distinct *SV2C* expressing L2-3 (*LAMP5^+^*) subtypes as In4a (*COL5A2^+^*), In4b (*COL5A2^-–^,EYA4^+^*) and In5 (*EYA4^+^ NOS1^+^*) (**Supplementary Figure 4**); layer 4/5 (*SULF1^+^*) located subtypes expressing *PVALB*, In6a (*RYR1^+^*)^6^ and the more peripheral and the most prevalent (8.65%) In neuron subtype in our cortical cell atlas In6b (*TAC1^+^*) ; and finally *SST* expressing L6 (*SYNPR^+^*) subtype In8 (*STXBP6^+^*) but not the other *SST -*positive In7 sub-population^6^ (**Supplementary** **Figure 4**).

**Data processing of published transcriptomic data for comparison with 10x RNAseq data**

***Laser- capture microscopy cell-types in the SN***

Microarray profiles of laser-captured human dopaminergic neuron from control and PD patients were obtained from GEO ([GSE24378](https://www.ncbi.nlm.nih.gov/geo/query/acc.cgi?acc=GSE24378))^8^. We downloaded the raw intensity values and performed a **Robust Multi-array Average** (**RMA**) transformation on probe-level with function RMA of R library oligo^10^, and computed the expression by gene by using the mean of expression probe measures, which were then log-transformed. Only the 9 control samples were used for the comparison.

Fastq files for RNA-seq data of **laser- capture microscopy** (**LCM**) astrocytes and microglia for a total of 18 samples for each cell-type in the SN^9^ were downloaded from **National Institute on Aging Genetics of Alzheimer's Disease Data Storage Site** (**NIAGADS**) ([NG00057](https://www.niagads.org/datasets/ng00057)). The paired-end sequencing data for each sample were mapped to the human reference transcriptome (GRCh38; Ensembl release 91) and quantified transcript abundance counts were obtained with the tool Salmon (v 0.11.0)^11^, using mapping based mode with default parameters, automatic library type inferring and additional options --seqBias,--posBias & --gcBias to account for sequence-specific biases, fragment -level GC biases and to account for 5’ or 3’ positional biases in the data. The transcript abundances were imported and summarized to gene-level using the R library tximport^12^. Only the protein-coding genes were used and data were filtered to include genes with >20 counts across all samples. The data were then normalised using the **variance stabilizing transformation** (**VST**) with vst function in the R library DESeq2^13^. Sample outliers were removed after visualising VST transformed data as PCA plots. Technical effects associated with the total number of genes were regressed out by using the removeBatchEffect function in the R library limma^14^. Finally, vst-transformed data for the control samples (n=9) for the LCM astrocytes and microglia were taken forward for comparison with the averaged log-transformed TPM 10x SN cell-type expression data. We discarded protein-coding genes that were not expressed in all three datasets(10x SN, LCM DaN & LCM astrocytes and microglia) used to perform these comparisons: 11,703 protein-coding genes were used in the comparisons for 28 samples/cells.

Observing that the distributions of gene expression levels were different between the datasets even after the ComBat batch correction of the R library sva^15^, we applied a non-parametric approach by ranking the genes according to their expression level in each tissue/cell. From this rank matrix, we computed pairwise spearman correlations between LCM cell-types/cell and performed clustering analyses by using Ward’s hierarchical approach and visualised the resulting matrix as a heatmap (**Supplementary Figure 13**).

***Single cell -RNA sequencing Temporal cortex***

For comparison with single-cell RNA-seq data from human **temporal cortex** (**TC**)^7^, we obtained gene count data from GEO ([GSE67835](https://www.ncbi.nlm.nih.gov/geo/query/acc.cgi?acc=GSE67835)) and converted these to TPM counts and restricted the analysis to protein-coding genes (n= 15,081) and cells (n= 465), maintaining the cell-type annotations from the original publication^7^. The expression matrix was then normalized and linearly regressed for batch identity with Seurat as described above, using a minimum cutoff of 1,000 protein-coding genes detected in at least three cells. HVG were identified from a mean variability plot (average expression versus dispersion (variance/mean) assigned to 20 bins based on average expression) using a log(variance/mean) cutoff of 0.5 to identify 3,662 genes. Moreover, differentially expressed genes associated with each cell-type (excluding the hybrid cell-type) were identified by Seurat, using parameters and thresholds as above. Transcriptional Pearson correlations were calculated in a pairwise manner across the averaged cell-type populations from the external and 10x datasets, grouped via Ward’s hierarchical clustering and visualised as a heatmap (**Supplementary Figure 13)**. The final comparison was restricted to log-transformed averaged cell-type TPM expression values of 6,446 genes, composed of HVG (protein-coding only) from both datasets along with the top 5 enriched genes (logFC > 2) in each cell-type.

***Single nuclei -DropSeq visual and frontal cortex***

We obtained the UMI counts for the **visual cortex** (**VC**) and **frontal cortex** (**FC**) single nuclei drop-seq data from GEO ([GSE97930](https://www.ncbi.nlm.nih.gov/geo/query/acc.cgi?acc=GSE97930)) and restricted the analysis to protein-coding genes (n= 10,919) and annotated nuclei (n= 29,441) maintaining cell-type annotations from the original publication^6^. As above, the expression matrix was then loaded into Seurat and nuclei with fewer than 300 UMI counts or more than 5000 UMI counts were omitted and further filtering to only retain nuclei with a minimum of 200 genes detected in at least 1% of cells was also, performed. We normalized and linearly regressed for technical effects associated UMI coverage and batch identity like the original publication with Seurat as above. HVGs were identified from a mean variability plot using a log(variance/mean) cutoff of 0.5 to identify 2,181 genes. Moreover, differentially expressed genes associated with each cell-type were identified by Seurat, using parameters and thresholds as above. Transcriptional Pearson correlations were calculated in a pairwise manner across the averaged log-transformed cell-type and subtype populations expression values from the external and 10x cortex dataset, grouped via Ward’s hierarchical clustering and visualised as a heatmap (**Supplementary Figure 13**). The analysis was restricted to 3,614 genes composed of HVG (protein-coding only) from both datasets and top 5 differentially expressed genes (logFC > 2) defining each cell-type.

**Cell-type association analysis**

***Cell-type association methods:*** To reduce the chance of spurious conclusions, we performed cell-type association analyses with two polygenic approaches**, stratified LD score regression** (**LDSC**) method^16^ and the **Multi-marker Analysis of GenoMic Annotation** (**MAGMA**) gene set analysis method^17^. The stratified LD score regression consists to partition heritability from genome-wide association studies. (GWAS) summary statistics (**Supplementary Table 4**) to different sets of genes specifically expressed in cell-type and to identify disease-relevant cell-type. MAGMA performs gene-set enrichment analysis based on GWAS summary statistics while accounting for LD structure between SNPs. A competitive gene-set analysis linear regression model was performed using MAGMA to test the hypothesis that a cell-type-specific gene set has a greater mean association with the complex trait than the genes not present in the gene set. A previous study comparing these two methods has shown that MAGMA can identify more significant results or false positives than LDSC, which the authors believe could be due to uncorrected genomic confounding that can be corrected by including gene-level covariates^16^. While both methods are based on different assumptions and algorithms and could have different results for certain traits, in this study, both methods include gene-level covariates such as gene size, gene sample size and linkage disequilibrium in the regression models to correct for confounding biases. We showed here that globally, there is agreement between the results given by both methods. Based on the Spearman rank correlations of cell-type association strength (-log_10_ Pvalue) between LDSC & MAGMA for each complex trait, we observed a very similar overall ranking of the cell-types across the SN (R=0.55, p<10^-16^) and the cortex (R= 0.53, p<10^-14^ ) (**Supplementary Figure 14**) for most brain-related complex traits such as schizophrenia, major depression disorder and bipolar disease. Moreover, non-brain-related traits (e.g. Height) also, show high positive correlations across the two methods.

***Cell-type-specific gene sets:*** To define cell-type-specific gene sets from the TPM expression matrix and the brain cell-types identified, we followed the approach of Finucane *et al.* 2018^16^. Here, for each gene a t-statistic is calculated between the expression of that gene in a given cell population as compared to its expression in all other cells, with cell-type-specific gene sets defined as the 10% of genes with the highest t-statistic in that cell-type (**Supplementary Data 4 and Supplementary Data 5**). Note that genes are not necessarily exclusive to a single cell-type. For the GTEx-brain tissue gene set, we used the GTEx brain gene set defined by Finucane et al. 2018 and downloaded the following file <https://data.broadinstitute.org/alkesgroup/LDSCORE/LDSC_SEG_ldscores/GTEx_brain_1000Gv3_ldscores.tgz>.

***Locus definition:*** On either side of the transcribed region of each gene in the set of cell-type specific genes, we extended the locus to define cell-(or tissue-) specific genomic region, aiming to capture regulatory elements that could affect the expression of that gene and mediate any GWAS variant effect on a trait. To define the size of this extension, we exploited the robust association between schizophrenia risk loci and genes specifically expressed in Ex neurons^18^. Thus, we ran LDSC ^16^ cell-type analysis with different windows size (5kb,10 kb, 25 kb, 50kb,75 kb and 100 kb) to evaluate the association between schizophrenia risk loci and Ex neuron gene expression and found that extending 25k around the transcribed region of each gene produced the most significant p-value for identifying Ex enrichment for schizophrenia (**Supplementary Figure 15**). We followed a similar approach for MAGMA^17^ and ran the gene-level based trait analysis with multiple window sizes for schizophrenia GWAS^19^, and evaluated the p-value for cell-type-specific association of cortical Ex and In neuron populations from our study with schizophrenia common genetic risk (**Supplementary Figure 16**). As with LDSC above, the most significant p-value for both Ex and In neurons was found to be 25 kb upstream and downstream of each gene and thus we used a 25kb window size for MAGMA analyses for all traits. Furthermore, we evaluated how the samples displaying amyloid angiopathy may affect the results of cell-type association analysis (individual 3, **Supplementary Table1**) and found consistent results in the LDSC association for all different traits with and without nuclei of this individual: **Supplementary Figure 17** (SN: R=0.82 (p <10^-16^) Cortex: R=0.93(p< p <10^-16^)).

***Conditional analysis for multiple cell-types within and across regions:*** Where LDSC identified multiple cell-types associated with the same trait, we used conditional analyses to evaluate whether it was the same genetic variants acting in different cell-types or distinct sets of genetic variants suggesting different cellular aetiologies. To test the association of cell-type A conditioned on cell-type B, we replaced the control gene set of all protein-coding genes expressed in both the cortex and nigral cell atlases by the cell-type B gene set and re-ran LDSC. When this conditional test produced a p-value associated with an LDSC Coefficient < 0.05, we considered that both cell-types could be involved independently in the same trait.

***Correction for the cell-heterogeneity between cortex and SN:*** In the case of traits for which genetic variants were simultaneously associated with SN and cortical cell-type, potential bias was that the different proportion of glial versus neuronal cell-types between these two brain regions may lead to the identification of different gene markers for the same cell-type (e.g. microglia) and suggested erroneously two distinct cell-type signals. To discard this bias for the conditional analysis, we created SN and cortex cellular atlas including the same number of astrocytes, microglia/OPC/ODC/neurons (398 Astrocytes, 72 DaNs or Ex neurons, 123 GABA neurons or In neurons, 325 microglia cells, 160 oligodendrocyte precursor cells and 147 oligodendrocytes cells) by random sampling cells in the original cellular atlas and performed the same LDSC analyses as with the original cellular atlases. As the endothelial cells were only identified in SN and as we found no evidence of co-association for the endothelial cells with a cortical cell-type for no trait, we discarded the endothelial cells in the creation of these homogenous cellular atlases. Globally, we noted a good agreement with LDSC done with the original cell atlas (**Supplementary Figure 6**) (SN R=0.83 (p < 10^-16^), cortex R=0.72 (p < 10^-16^)), nevertheless we noted a p-value inflation in the microglia cells identified in the cortex for SCZ and bipolar disorder. We then repeated the conditional LDSC analysis for traits with SN and cortical cell-type enrichment (**Supplementary Table 8**, column P-value (LDSC) (Matched Cellular Atlas)).

An RMarkdown document, including a version with R code to generate these gene sets and perform cell-type association, can be found here: https://github.com/csandorfr/SN_Atlas.

**Functional Analysis - Protein Protein Interaction network**

A combined protein-protein interaction network was created based on diverse resources: BioGRID 3.4 (accessed on September 2017)^20^, HitPredict (accessed on September 2017)^21^, IntAct (accessed on September 2017)^22^, STRING (accessed on September 2017, restricted to Homo sapiens and experimental scores higher than zero) ^23^, CORUM (accessed on September 2017)^24^and Reactome (accessed on September 2017)^25^. The combined network consisted of a total of 20,591 genes and 1,973,967 interactions.

**Work Cited**

**Supplementary Figures**

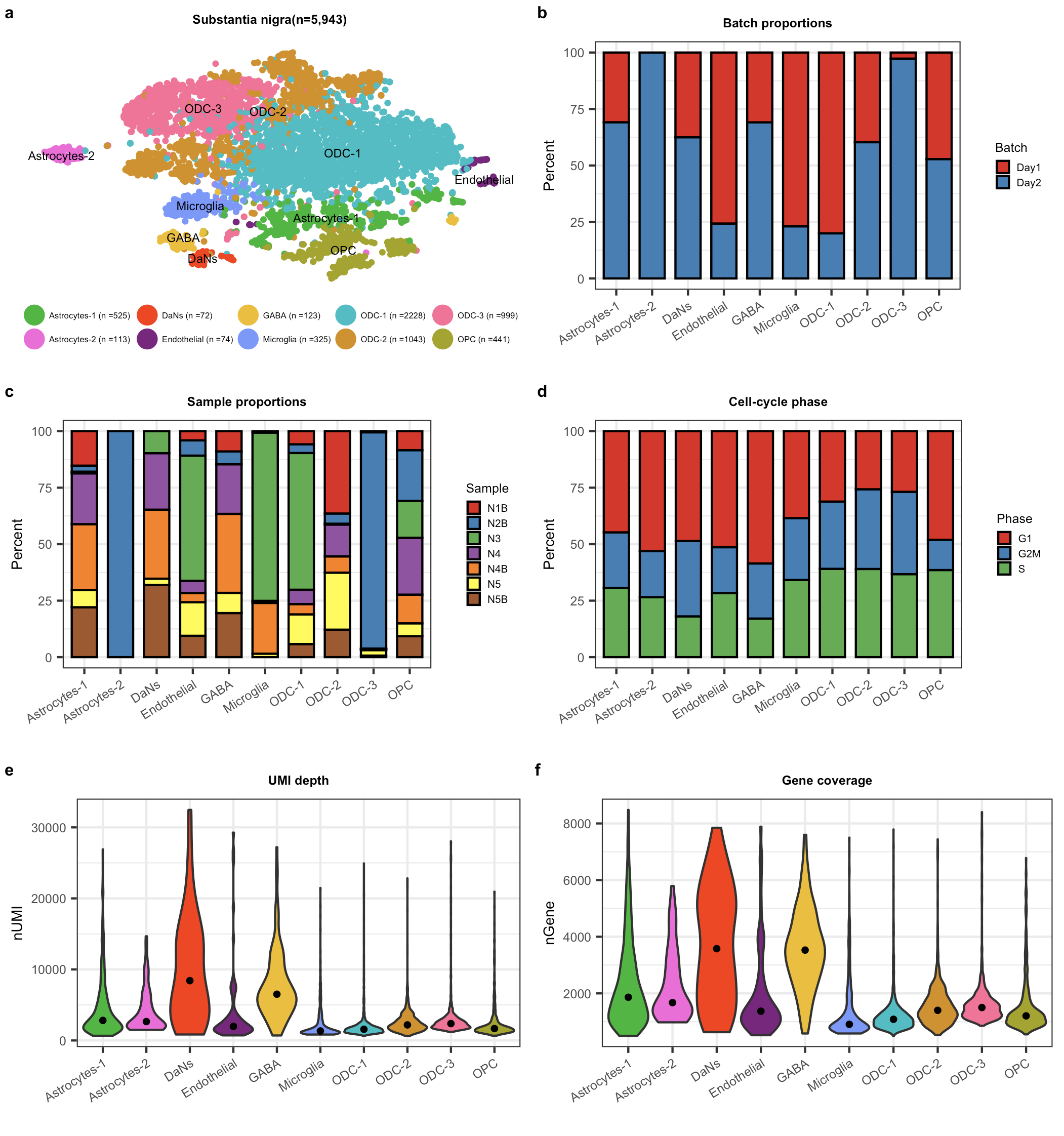

**Figure S1: Covariates distribution in the substantia nigra (SN) cell atlas.**

a) T-distributed stochastic neighbor embedding (tSNE) plot for all non-neuronal (astrocytes, endothelial, microglia, ODC and OPC) and neuronal (DaNs and GABA) cell-types or subtypes across the SN. b) barplot plot showing batch proportions per level-2 cell-type. c) barplot as in (b) showing sample proportions per level-2 cell-type. d) barplot as in (b) showing cell cycle phase proportions across level-2 cell-types. e) Violin plot showing unique molecular identifier (UMI) depth across all level-2 cell-types. f) Violin plot as in (e) showing gene coverage across all level-2 cell-types. In (e) and (f) the black dot denotes the median UMI and genes for the cell-type. DaNs:Dopaminergic neurons; GABA: GABAergic neurons; ODC: Oligodendrocytes; OPC: Oligo-precursor-cells.

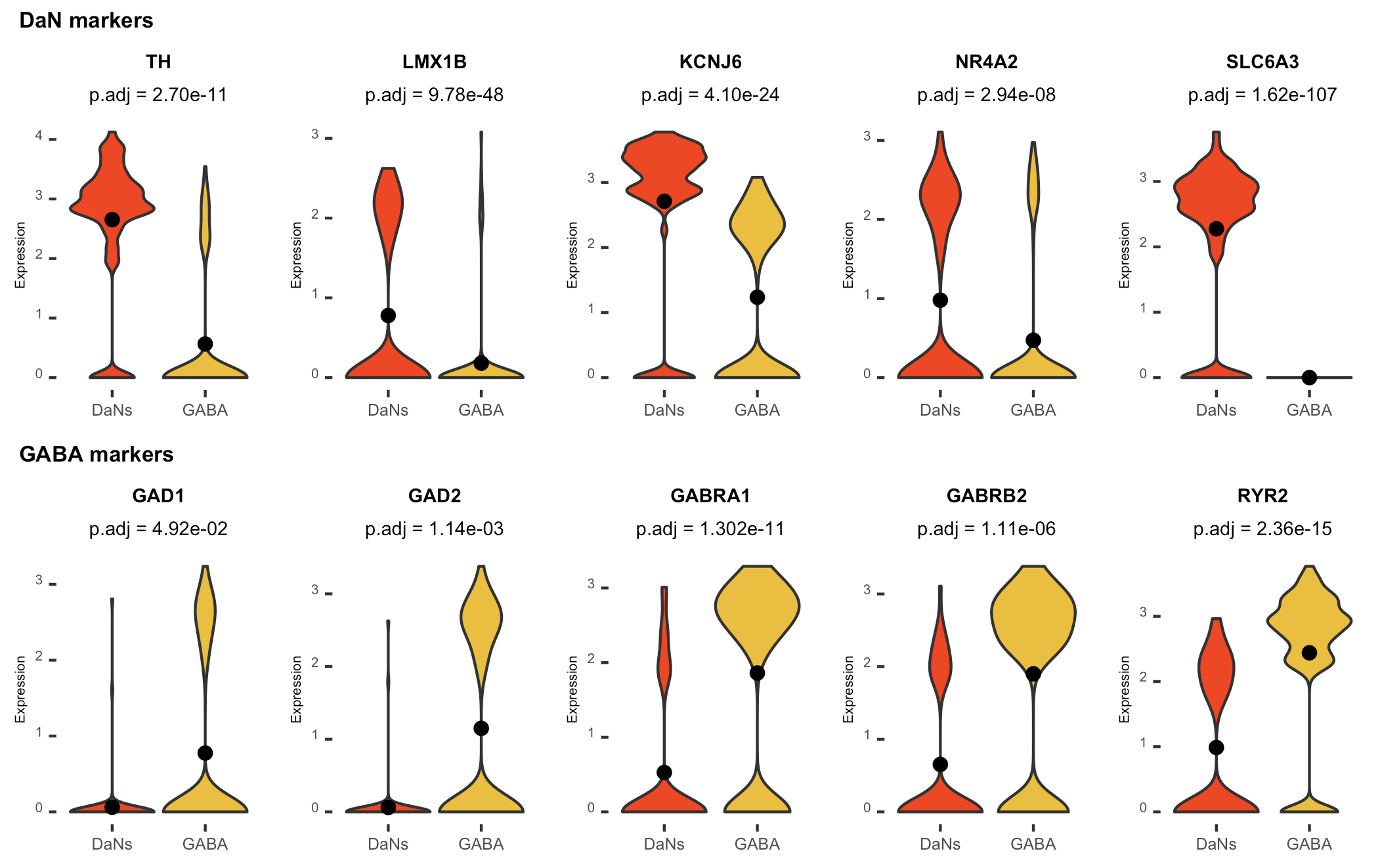

**Figure S2: Violin plots of DaN and GABA neuron markers in the SN neurons.**

Violin plots of expression values (log10 TPM values) for known DaN and GABA markers. Differential expression p-values after FDR correction(p.adj) using the negative binomial test in Seurat were calculated between the two neuronal populations (Methods & data S3). In each of the violin plots the black dot denotes the mean expression for the marker genes for the cell-type. DaNs: Dopaminergic neurons; GABA: GABAergic neurons.

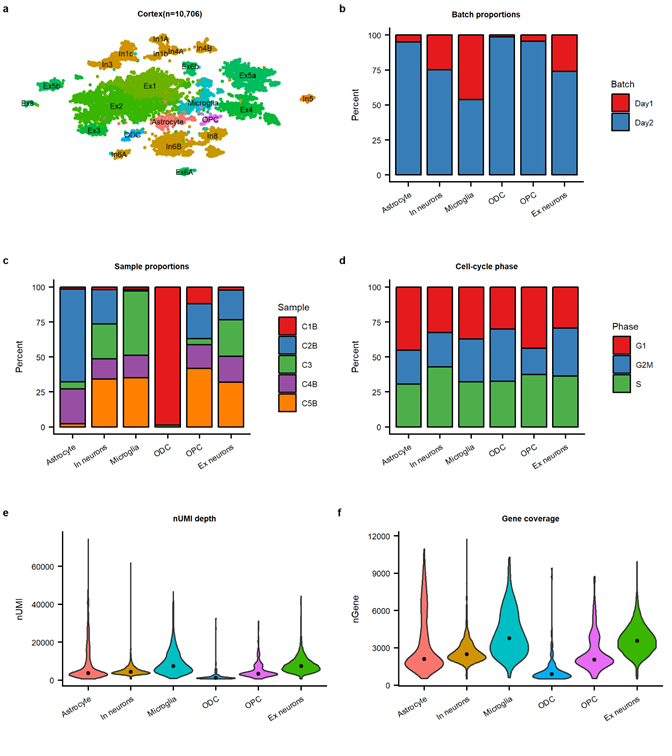

**Figure S3: Covariates distributions in the cortex cell atlas.**

**a)** T-distributed stochastic neighbor embedding (**tSNE**) plot for all non-neuronal (astrocytes, microglia, ODC and OPC), Ex and In neuron cell-types or subtypes across the cortex. **b)** barplot plot showing batch proportions per level-1 cell-type. **c)** barplot as in **(b)** showing sample proportions per level-1 cell-type. **d)** barplot as in **(b)** showing cell cycle phase proportions across level-11 cell-types**. e)** Violin plot showing unique molecular identifier (**UMI**) depth across all level-1 cell-types**. f)** Violin plot as in **(e)** showing gene coverage across all level-1 cell-types. In **(e)** and **(f)** the black dot denotes the median UMI and genes for the cell-type. Ex:Excitatory;In:Inhibitory ;ODC: Oligodendrocytes;OPC: Oligo-precursor-cells.

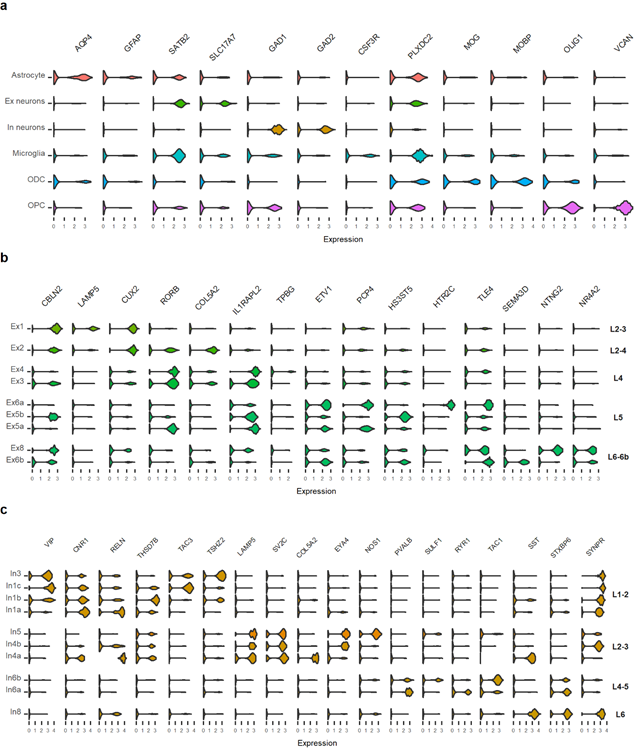

**Figure S4: Cortex cell-types:**

(a) Violin plots of expression values (log10 TPM values) of cell-type specific markers for the cell-types in the cortex. (b) Violin plots of expression values (log10 TPM values) of cell-type and cortical layer-specific enriched marker gene profiles^6^ for the Excitatory neurons subtypes. (c) Violin plots of expression values (log10 TPM values) of cell-type and cortical layer-specific enriched marker gene profiles for the Inhibitory neuron subtypes. Ex: Excitatory; In: Inhibitory, ODC: Oligodendrocytes; OPC:Oligo-precursor cells, L1-2; Layer 1/2, L2-3; Layer 2/3,Layer 2-4; Layer 2/3/4, L2 ; Layer2, L3; Layer 3, L4; Layer 4, L4-5; Layer 4/5, L6; Layer 6; Layer 6-6b; Layer 6/6b

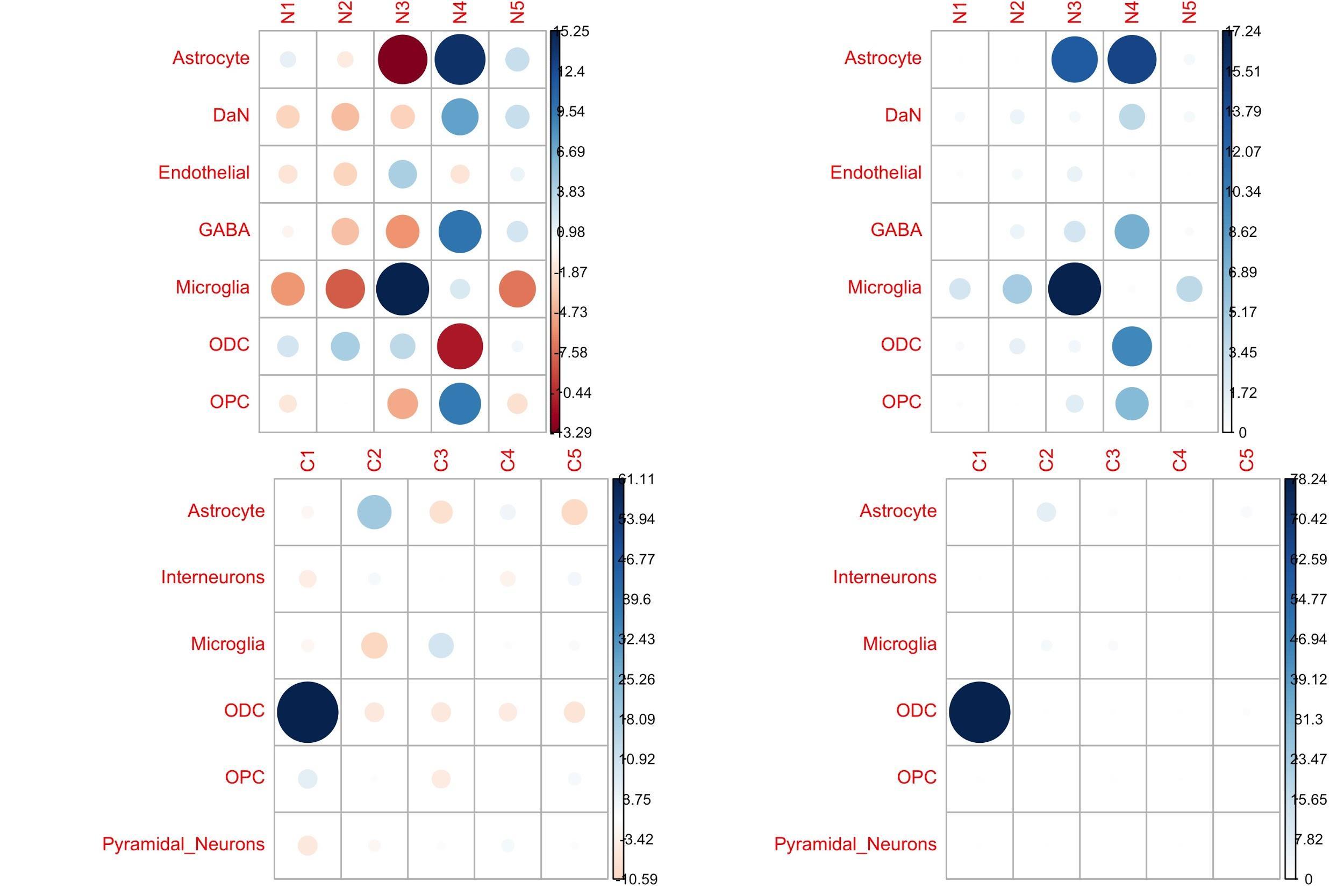

**Figure S5: Only the minor cell populations show difference in the frequency between the individuals.**

Cell composition (left) & contribution (right) of each individual within SN (top) and cortex (bottom) single nuclei atlas. (Left graph) within each atlas, we calculated the Pearson residual reflecting an under (red) or over-representation (blue) of a specific cell-type for an individual. (Right graph) we calculated the % represented by chi2 residual reflecting whether an individual contributes more to a cell-family category than another individual.

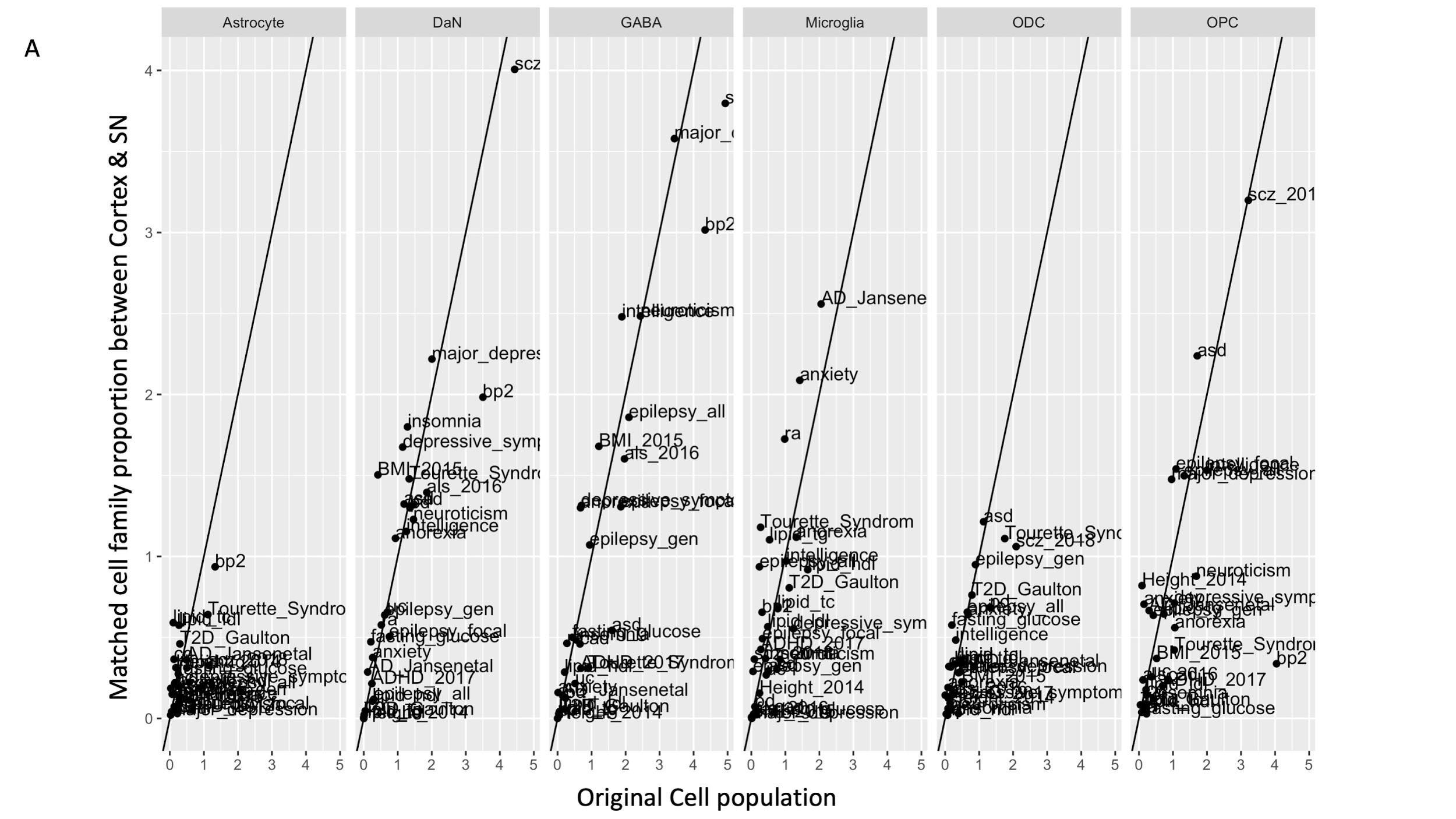

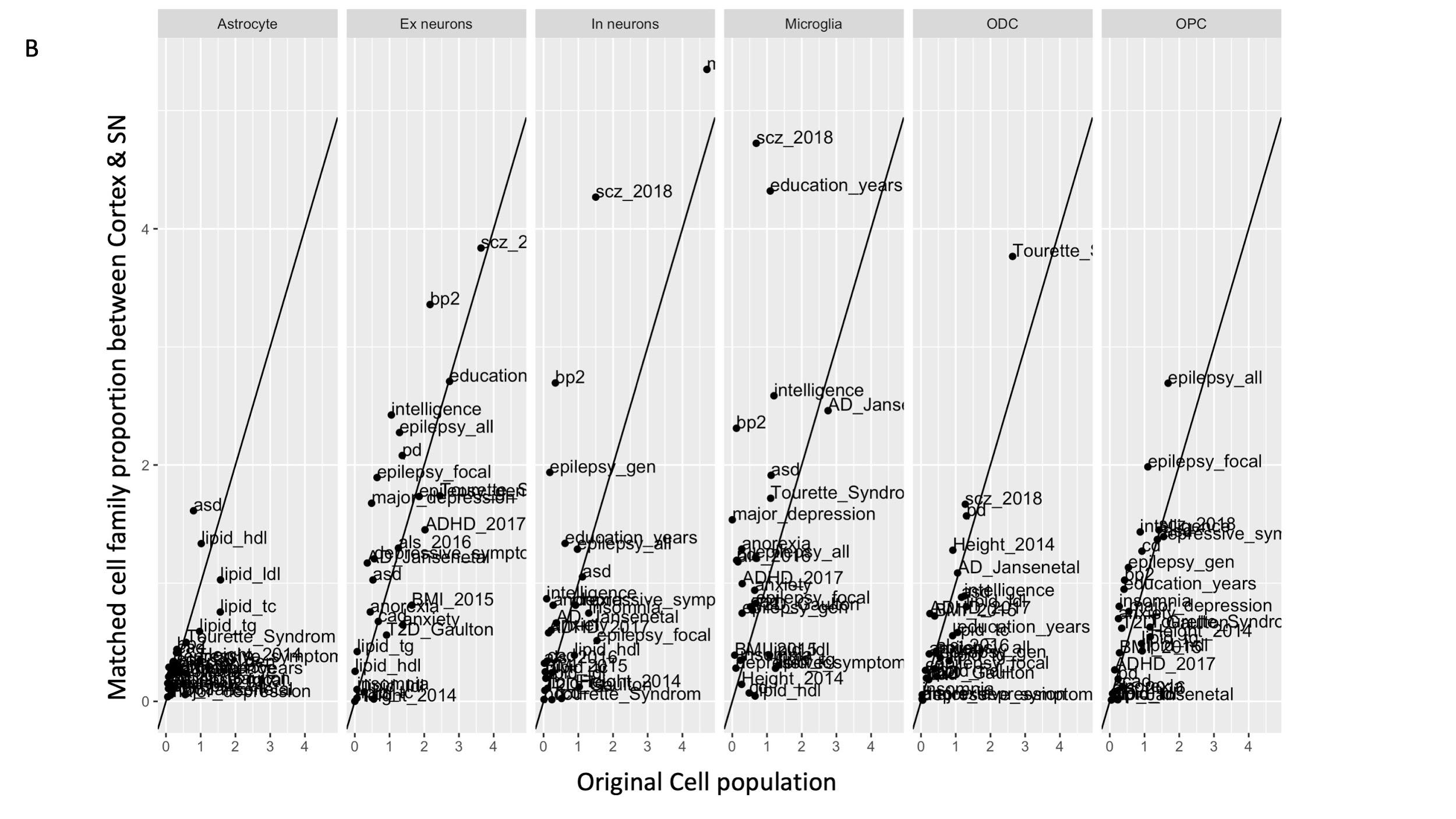

**Figure S6: Comparison of cell-type enrichment analysis performed with LDSC for different traits between the cellular atlases with matched cell family proportion between cortex and SN and the original cellular atlases**

From the original cellular atlas, we created SN (A) and cortex (B) cellular atlas including the same number of astrocytes microglia/OPC/ODC/Neurons by random sampling cells in the original cellular atlas and performed the same LDSC analyses as with original cellular atlases (Methods). For a specific cell-type, the x-axis and y-axis of each plot represent the -log 10 p-value for enrichment of genetic variants associated with human trait by using the original cellular atlases and the matched cell family proportion cellular atlases respectively.

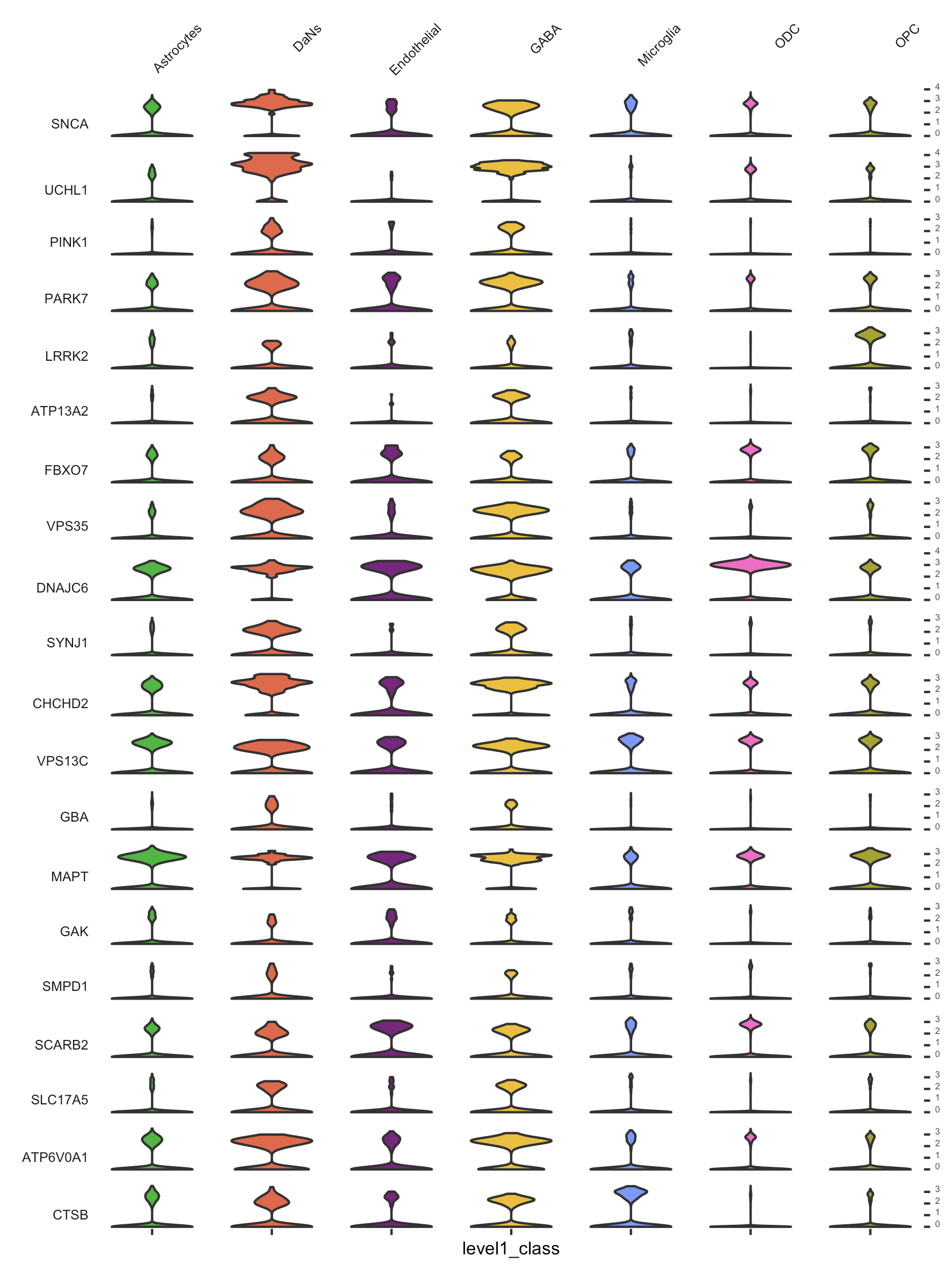

**Figure S7: Expression of known Parkinson’s disease genes in SN level-1 cell-types.**

Violin plots of expression values (log10 TPM values) of Parkinson's Disease (PD) risk genes^26^ for the cell-types in the SN.

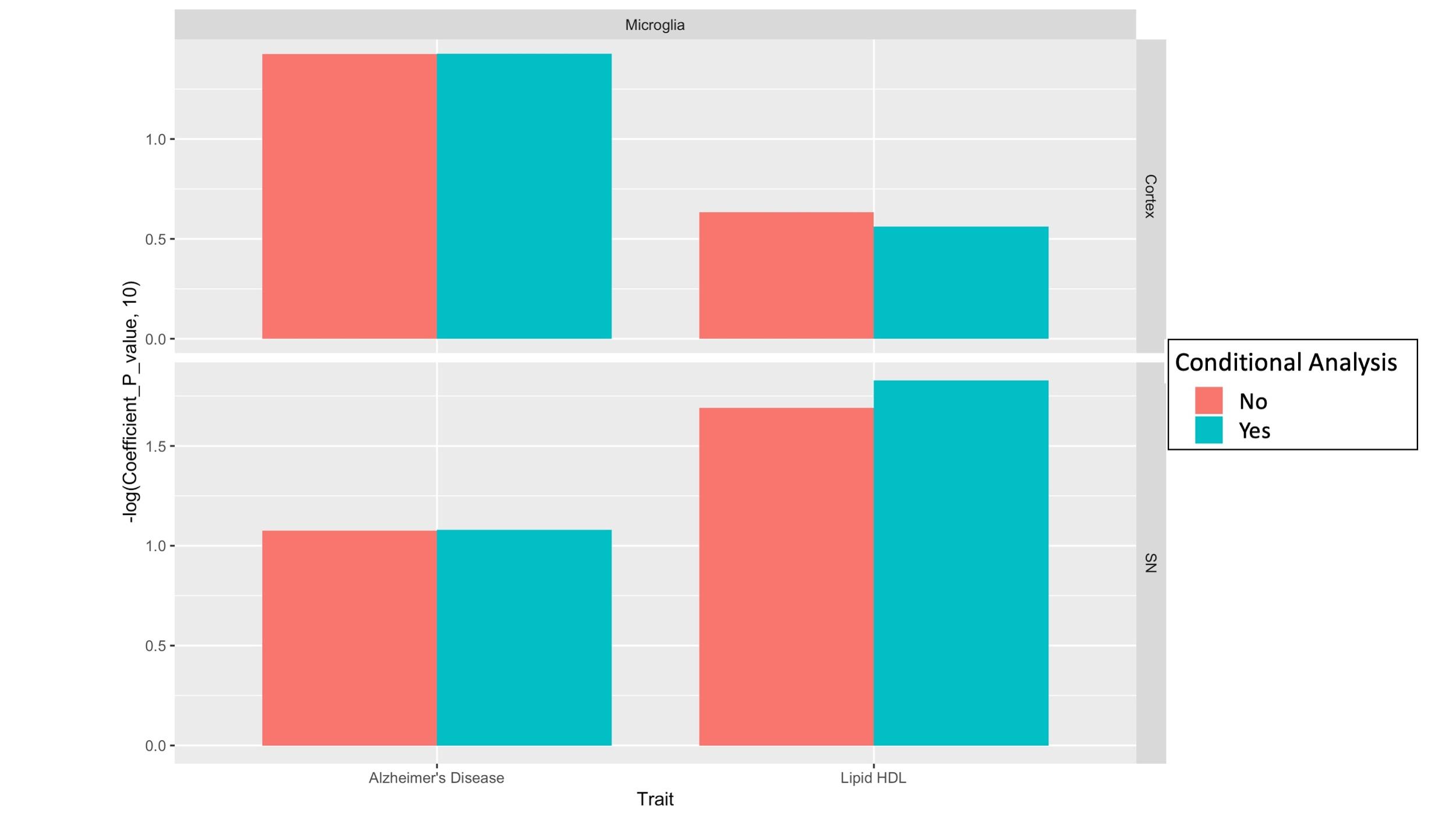

**Figure S8: The microglia specific signals for the genetic variants associated with HDL cholesterol level and AD correspond to distinct genetic aetiologies.**

LDSC analysis by using the AD and HDL GWA summary statistic adjusted (green) or not (red) for HLD and AD risk effect respectively to evaluate the cell-type association within microglia in the cortex (top) and SN (bottom). Each bar represents the -log 10 p-value of enrichment genetic variants associated with AD or HDL (adjusted or not for the other trait) for microglia cell-type identified in the cortex or SN.

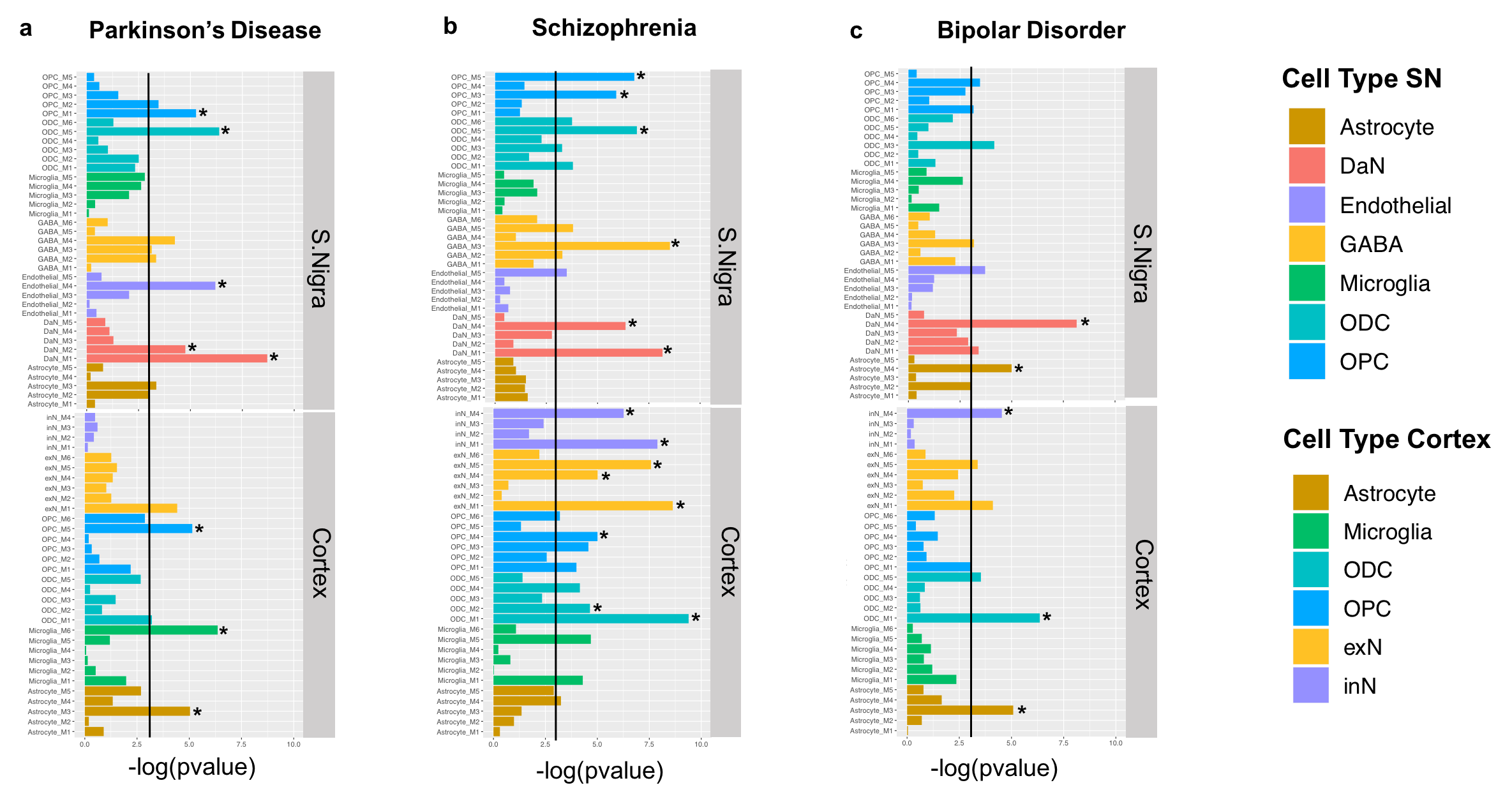

**Figure S 9: Convergence of PD-risk (a), SCZ-risk (b) and BP-risk (c ) across SN and cortex cell-type specific gene modules.**

We assessed SCZ-, PD- and BP- risk enrichment in cell-type specific gene modules from cortex and SN through MAGMA gene set analysis. Barplots represent -log_10_ p-values of enrichment, vertical lines correspond to p-value=0.05 and star symbols (*) indicate significance after multiple test correction. Only the associations to modules within cell-types that showed significance from the more general analysis (Figure 2) were discussed further.

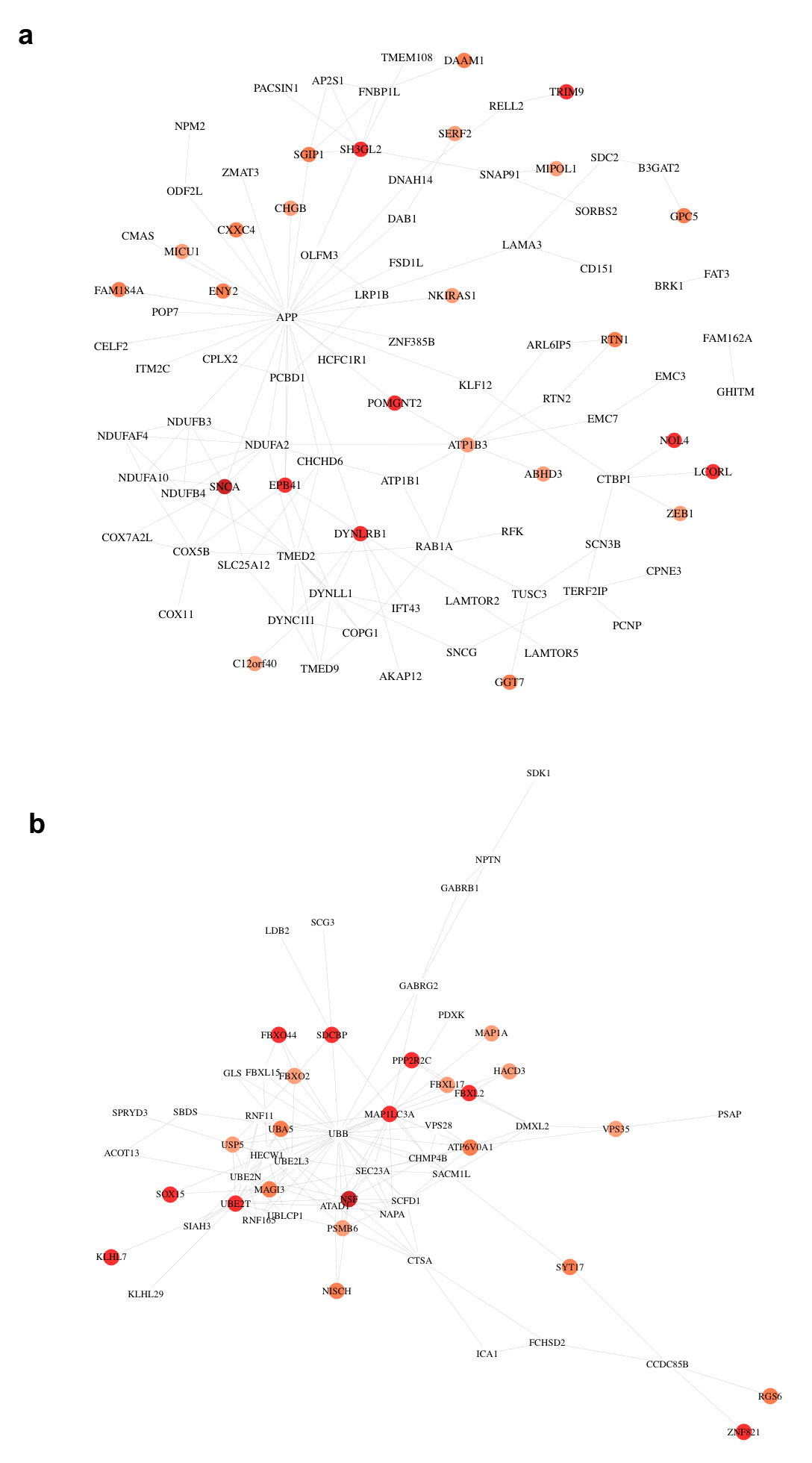

**Figure S10: Network representation of PD-risk genes within SN module DaN_M1**

**(a) and DaN_M2 (b).** Gradient of colours is proportional to PD-risk association determined by MAGMA adjusted_zscores, with darker colours associated to higher adjusted_zscores (higher disease-risk association). White nodes correspond to genes within the modules that link PD-risk associated genes together but not found directly associated to PD.

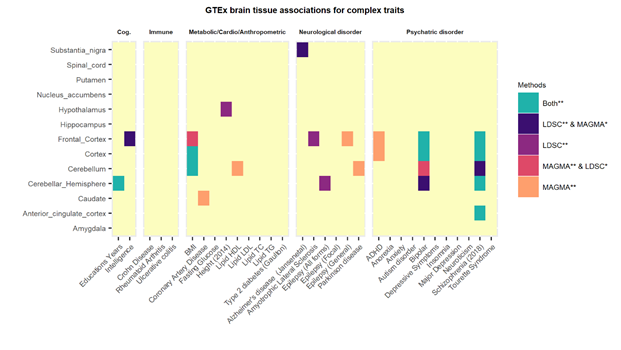

**Figure S11: Tissue- trait associations in diverse human complex traits by using GTEx bulk brain tissue transcriptomic data.**

Two classical approaches: stratified LD score regression (LDSC) and the MAGMA gene set analysis have been used to identify the association between multiple complex traits genome wide association signals within 13 brain regions from bulk GTEx transcriptomic data. The heatmap colour indicates if the p-value of cell-type enrichment for a trait is significantly associated with both methods or either methods after Bonferroni multiple test correction(**) and also, if nominally significant (p<=0.05) in the other method (*). Bonferroni threshold was P-value/No.of.cell-types. The different traits have been clustered by category: Cognitive phenotype (Cog.), Autoimmune disease (Immune), Metabolic, Cardiovascular and Anthropometric Traits (Metabolic/Cardio/Anthropometric) Neurological disorder, Psychiatric disorder.

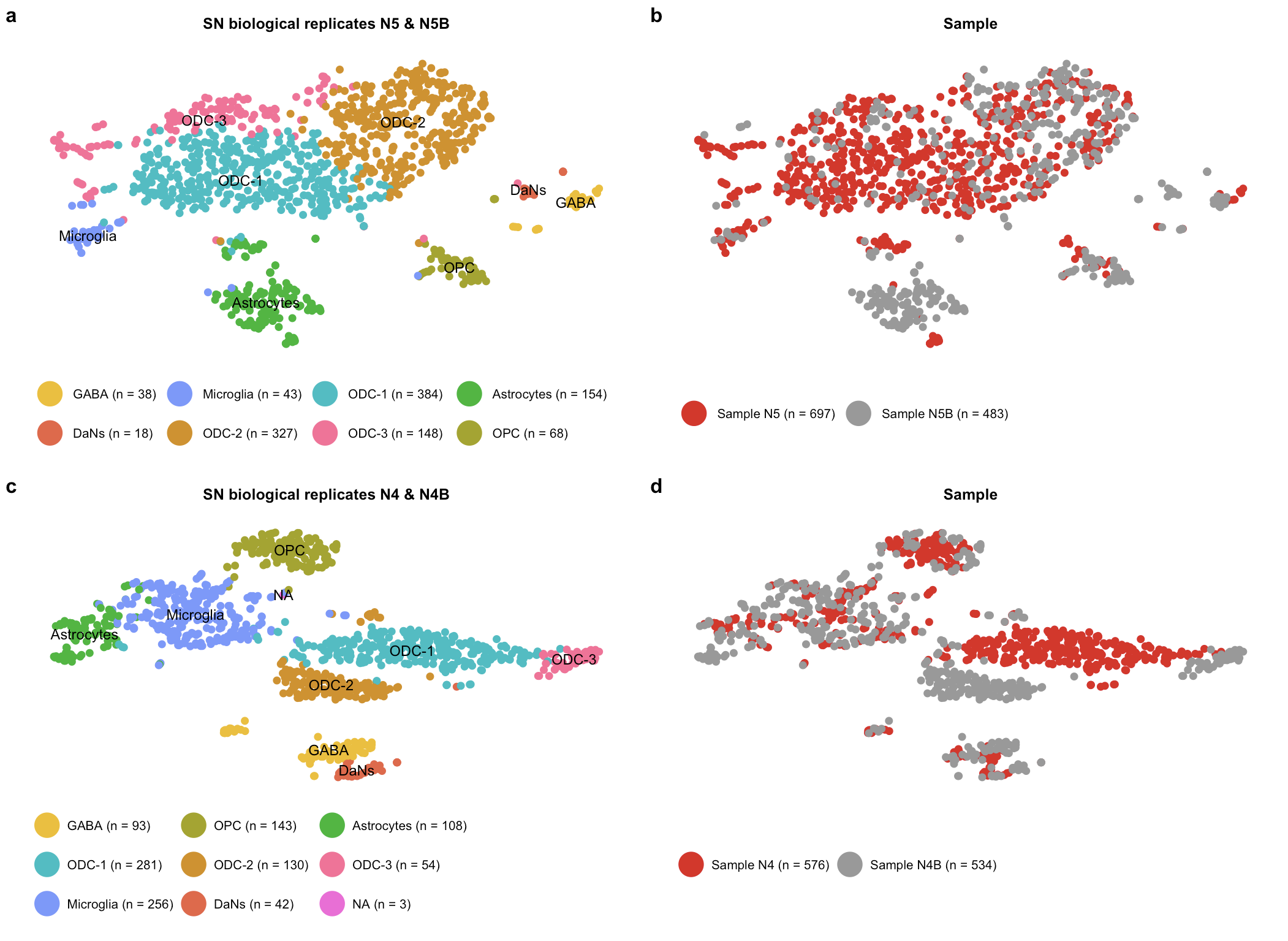

**Figure S12: Evaluation of similarity of substantia nigra (SN) biological replicates from the same individuals.**

a) T-distributed stochastic neighbor embedding (tSNE) plot for all non-neuronal (astrocytes, microglia, ODC and OPC) and neuronal (DaNs and GABA) cell-types or subtypes across biological replicates N5 and N5B in the SN. b) t-SNE plot as in (a) showing sample identity N5 and N5B (**Supplementary Table 1**). c) t-SNE as in (a) showing cell-types for N4 and N4B SN biological replicates (**Supplementary Table 1**). d) t-SNE as in (c) showing sample identity N4 and N4B (**Supplementary Table 1**). All major cell-types were identified in both sets of replicates from each individual. In both sets of replicates a distinct cluster of endothelial cell-type was not identified and in N5&N5B a small cluster of N=3 was not attributed to any particular cell-type. DaNs: Dopaminergic neurons; GABA: GABAergic neurons; ODC: Oligodendrocytes; OPC: Oligo-precursor-cells ; N4: Nigra 4;N4B;Nigra 4B;N5:Nigra 5; N5B: Nigra 5B.

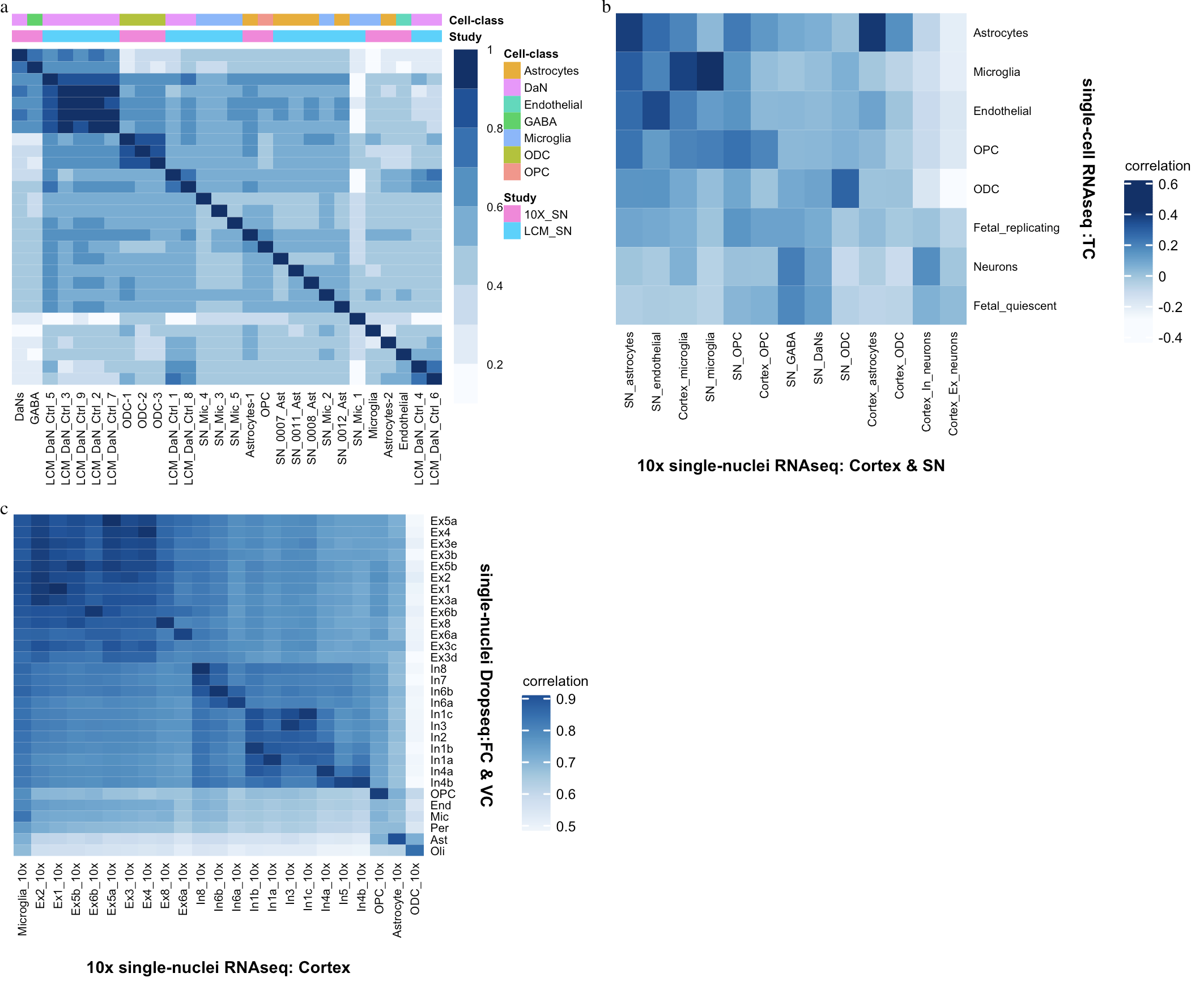

**Figure S13: Comparison of substantia nigra and cortex (10x) cell-types with public transcriptomic datasets**

a) SN laser- capture microscopy (LCM) Cell-types. A correlation heatmap comparing averaged SN level-2 cell-type populations with LCM SN Dopaminergic neurons (LCM_DaN_Ctrl)^8^, microglia (Ctrl_Mic) and astrocytes (Ctrl_Ast)^9^ from control samples. b) Cortex single cell RNA-seq data. Pairwise correlation comparison of the averaged 10x cortex and SN neuronal and non-neuronal subtype populations with the averaged temporal cortex (TC) single cell RNA-seq type populations ^7^. c) Cortex single nuclei Drop-seq data. Pairwise correlation comparison of the averaged 10x cortex neuronal and non -neuronal subtype populations with the averaged cell-type populations in the frontal cortex (FC) and visual cortex (VC) of this single nuclei dataset^6^.

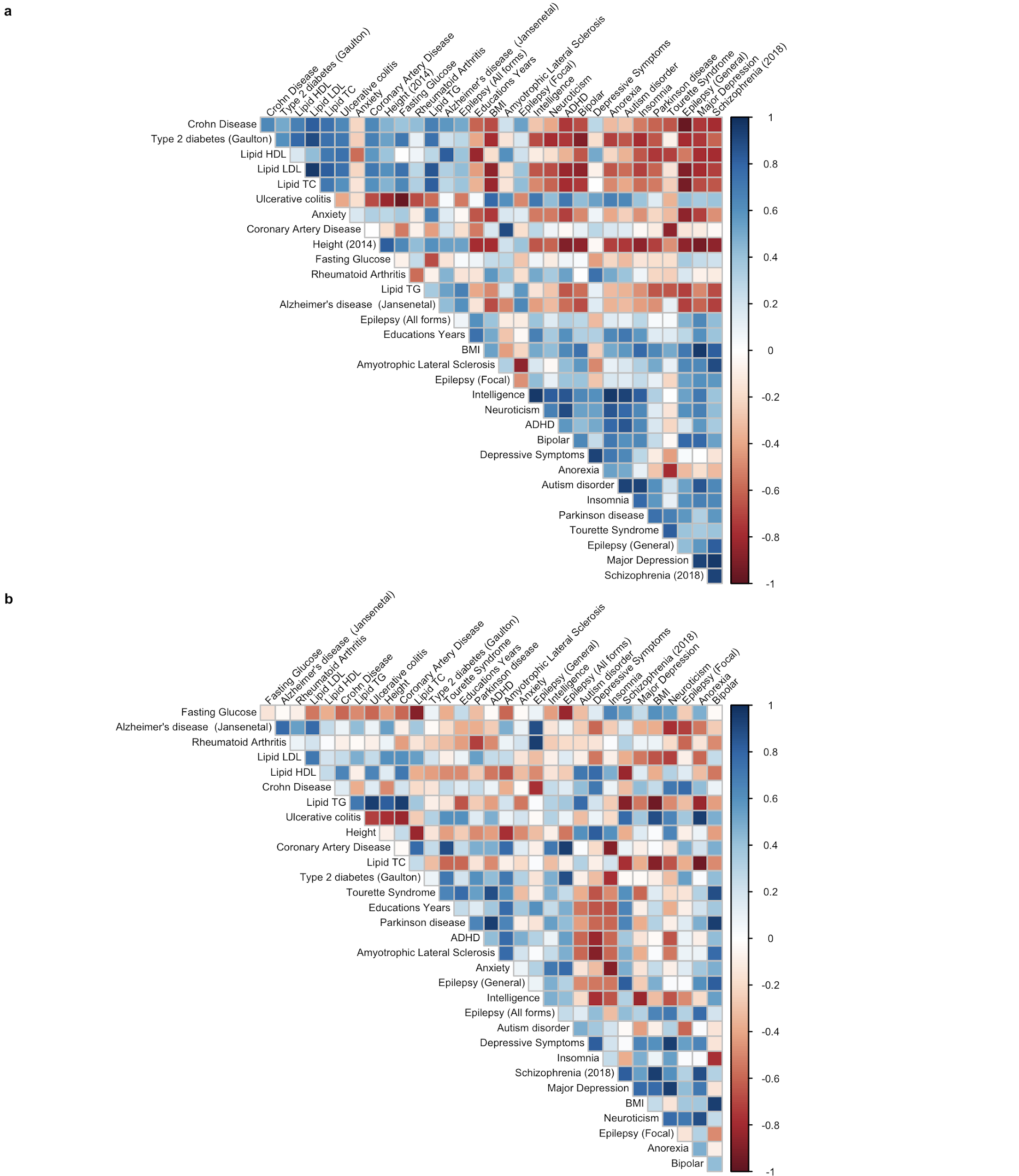

**Figure S14: LDSC vs MAGMA spearman rank correlation plot for cell-type associations (-log10 pvalues) in the SN and cortex cell atlas.**

The clustered Spearman rank correlations between the rank orders of cell-type associations(-log10Pvalue) for LDSC and MAGMA across traits for the a) SN and b) cortex cell-types are shown. A moderate to high positive correlation was observed for neurological disorders such as PD and ALS, for psychiatric disorders such as Schizophrenia, ADHD, Autism Disorder, Major Depression and Insomnia. Moreover non-brain metabolic traits such as Height, BMI and Cholesterol (HDL & LDL) and immune traits (Crohn's disease) were also, highly concordant between the two methods.

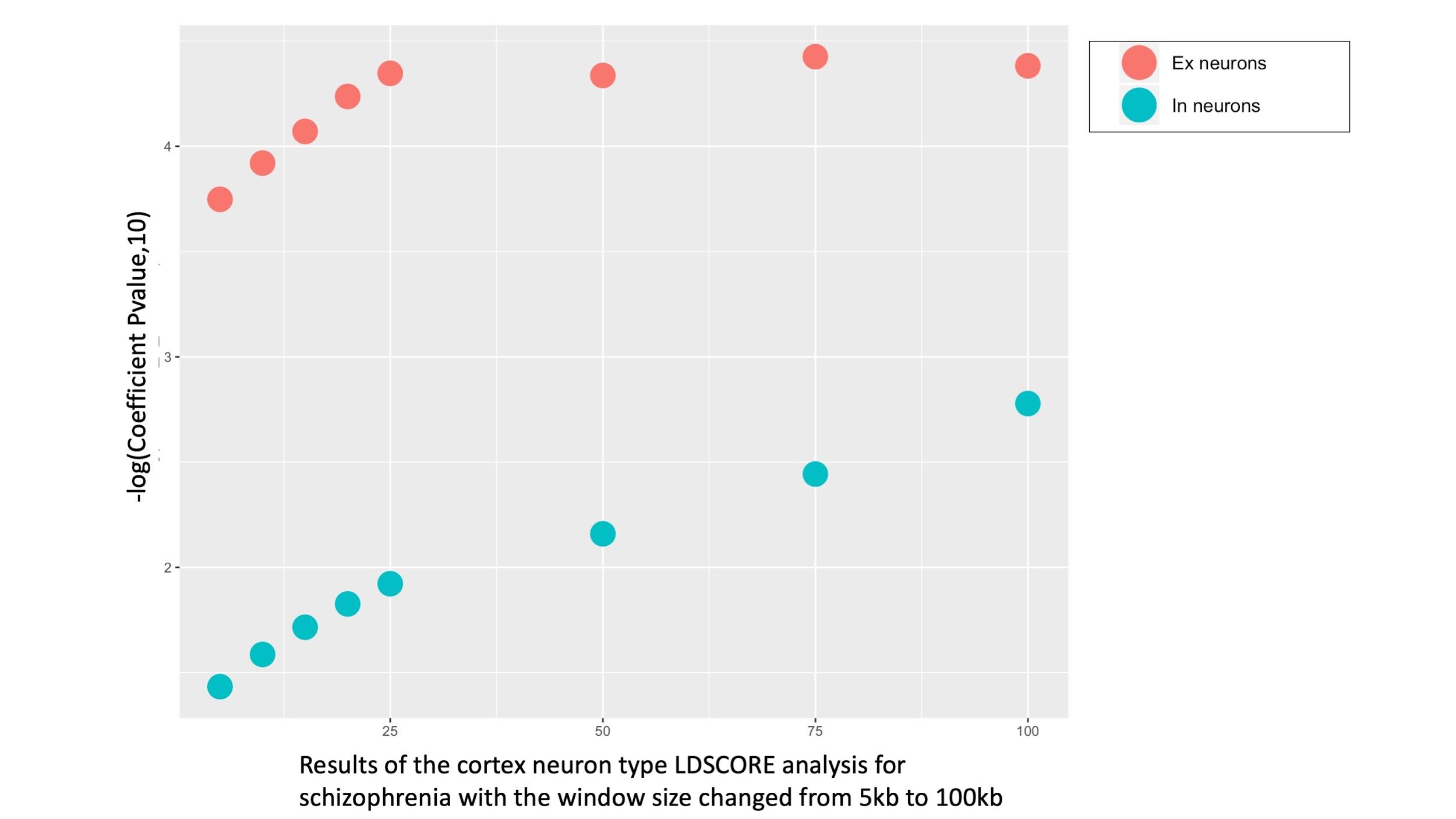

**Figure S15: Selection of windows size parameter around each transcribed region in the LDSC cell-type analysis.**

Results of the LDSC cell-type analysis with different windows size (5kb, 10 kb, 25 kb, 50kb,75 kb and 100 kb) to evaluate the association between schizophrenia and Excitatory neurons and Inhibitory neurons in the cortex. The x-axis represents the windows size added on either side of each gene, while the y-axis represents -log10 p-value associated with LDSC Coefficient.

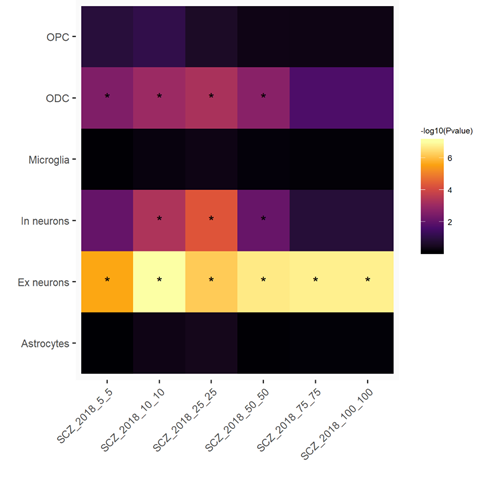

**Figure S16: Selection of window size parameter around each gene region for the MAGMA gene set based cell-type analysis.**

Heatmap showing the -log_10_ p value of the cell-type association analysis for Schizophrenia for the adult cortex single nuclei cell-types identified in this study at different incremental window sizes (5kb,10kb,25kb,50kb,75kb & 100kb). The window size was benchmarked on the schizophrenia p-value for Excitatory neurons (Ex) and Inhibitory neurons (In). Each cell has been labelled with an * if the -log10 p-value is >= to Bonferroni significance threshold (-log_10_( 0.05/No of Cell-types)). SCZ: Schizophrenia.

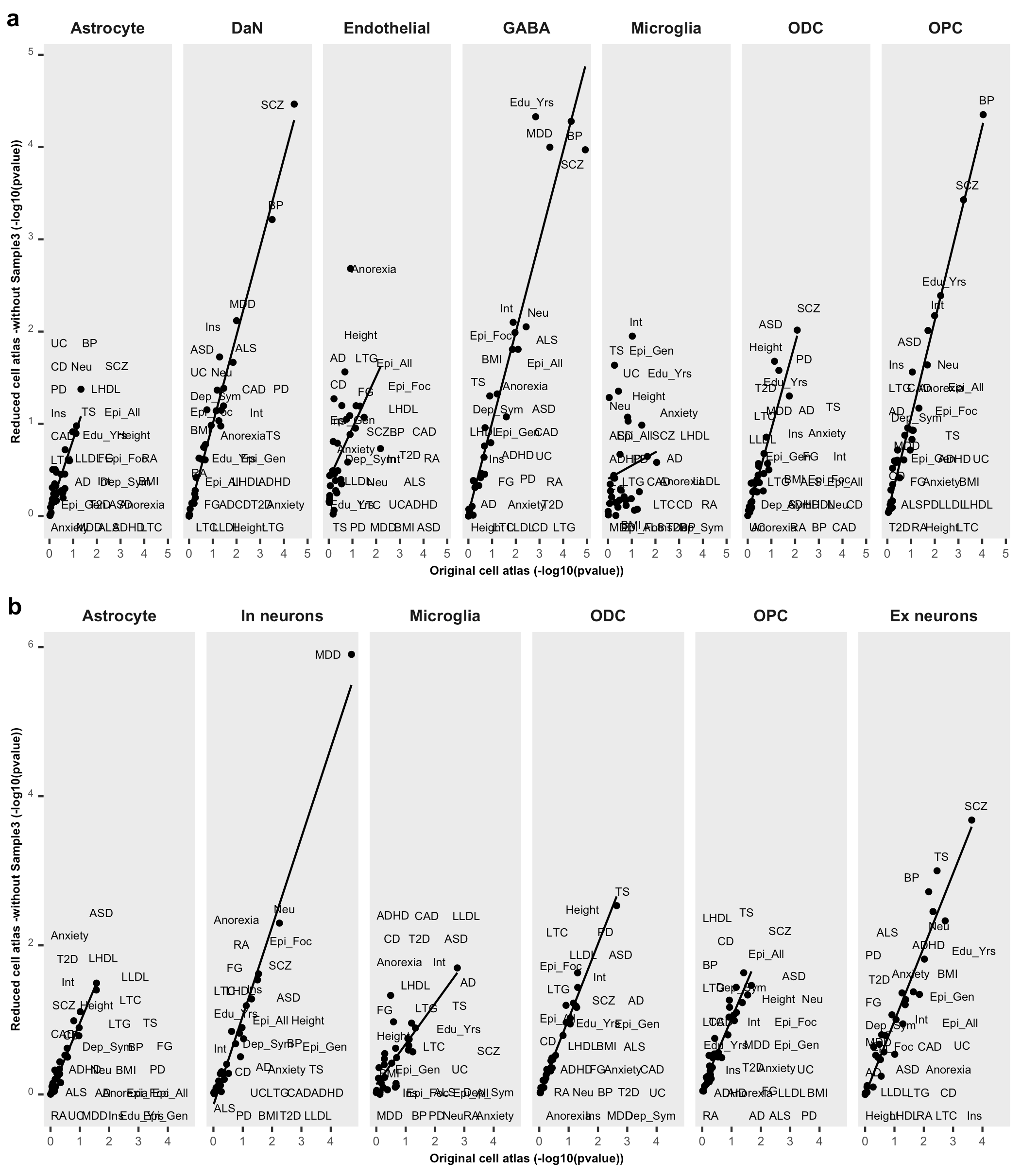

**Figure S 17: Comparison of cell-type enrichment (-log10 pvalues) for different traits between the original cell atlas and the reduced cell atlas (no Sample 3).**

The plot compares the cell-type association enrichments (-log10(pvalue)) between the cell atlases with and without the individual with amyloidosis (Sample 3, Supplementary Table1) in the a) SN and b) cortex with the LDSC method in each cell-type. For a specific cell-type, the x-axis and y-axis of each plot represent the -log 10 p-value of enrichment genetic variants associated with human trait by using the original cellular atlases and the reduce cell cellular atlases respectively.

**Supplementary Table Legends**

Supplementary Table 1: Single- nuclei RNAsequencing experimental design and sample information

Supplementary Table 2: Gene Ontology pathway analysis to distinct two sub-population of astrocytes in the Substantia Nigra

Supplementary Table 3: Substantia nigra and cortex cell population proportions

Supplementary Table 4: Source of genome-wide association studies meta-analysis summary statistics used in this study

Supplementary Table 5 : Risk association p-value for 30 brain and non brain-related traits with cell-types from the Substantia nigra (SN) using LDSC & MAGMA

Supplementary Table 6: Conditional cell-type analysis to evaluate whether multiple significant Substantia Nigra cell-type signature associated with the same trait coincide or not with the same genetic variants.

Supplementary Table 7: Risk association p-value for 30 brain and non brain-related traits with cell-types from the Cortex using LDSC & MAGMA

Supplementary Table 8: Conditional cell-type analysis to evaluate whether a significant Microglia/OPC/ODC/Neurons detected simultaneously in the SN & cortex with the same trait coincide or not with the same genetic variants.

Supplementary Table 9: Risk association p-value for 30 brain and non brain-related traits with GTEx brain region tissue using LDSC & MAGMA.

Supplementary Table 10: Top 100 GO enriched pathways for cell-type specific modules in substantia nigra

Supplementary Table 11: Top 100 GO enriched pathways for cell-type specific modules in cortex

**Supplementary Data legends**

Supplementary Data 1: 10x genomics single nuclei- RNA sequencing quality control metrics for samples

Supplementary Data 2: Cortex-substantia nigra cell-type annotations

Supplementary Data 3: Differentially expressed genes for cell-types and cell-type subpopulation clusters in the substantia nigra (SN) and cortex sn - RNAseq

Supplementary Data 4: Cell-type specific gene sets generated using t-statistics for cortex and SN cell atlas

Supplementary Data 5: Cell-type specific gene sets generated using t-statistics for cortex and SN cell atlas without the amyloidosis sample
